# Structural foundations of potassium selectivity in channelrhodopsins

**DOI:** 10.1101/2022.09.26.509509

**Authors:** Elena G. Govorunova, Oleg A. Sineshchekov, Leonid S. Brown, Ana-Nicoleta Bondar, John L. Spudich

## Abstract

Kalium channelrhodopsins (KCRs) are light-gated K^+^ channels recently found in the stramenopile protist *Hyphochytrium catenoides*. When expressed in neurons, KCRs enable high-precision optical inhibition of spiking (optogenetic silencing). KCRs are capable of discriminating K^+^ from Na^+^ without the conventional K^+^-selectivity filter found in classical K^+^ channels. The genome of *H. catenoides* also encodes a third paralog that is more permeable for Na^+^ than for K^+^. To identify structural motifs responsible for the unusual K^+^ selectivity of KCRs, we systematically analyzed a series of chimeras and mutants of this protein. We found that mutations of three critical residues in the paralog convert its Na^+^ selective channel into a K^+^ selective one. Our characterization of homologous proteins from other protists (*Colponema vietnamica, Cafeteria burkhardae* and *Chromera velia*) and metagenomic samples confirmed the importance of these residues for K^+^ selectivity. We also show that Trp102 and Asp116, conserved in all three *H. catenoides* orthologs, are necessary, although not sufficient, for K^+^ selectivity. Our results provide the foundation for further engineering of KCRs for optogenetic needs.

**IMPORTANCE:** Recently discovered microbial light-gated ion channels (channelrhodopsins) with a higher permeability for K^+^ than for Na^+^ (kalium channelrhodopsins, or KCRs) demonstrate an alternative K^+^ selectivity mechanism, unrelated to well-characterized “selectivity filters” of voltage- and ligand-gated K^+^ channels. KCRs can be used for optogenetic inhibition of neuronal firing, and potentially for the development of gene therapies to treat neurological and cardiovascular disorders. In this study we identify structural motifs that determine the K^+^ selectivity of KCRs that provide the foundation for that provide the foundation for elucidating their K^+^ selectivity mechanism and for their further engineering as optogenetic tools.

## INTRODUCTION

Channelrhodopsins (ChRs) are a diverse group of >500 light-gated ion channels found in eukaryotic microbes (1) and widely used as optogenetic tools (2). ChRs are members of a larger protein family known as microbial rhodopsins (3-5), and are composed of seven transmembrane helices (TM1-TM7) with the retinal chromophore attached in a Schiff base linkage to a conserved Lys residue in the middle of TM7. In the model flagellate alga *Chlamydomonas reinhardtii*, the role of ChRs as phototaxis receptors has been established by analysis of knockdown genetic transformants (6). ChRs are thought to function similarly in other microorganisms, because all species in the genomes in which ChRs have been found produce flagellate gametes and/or zoospores during their life cycles.

By their ion selectivity ChRs can be classified into three groups: ACRs, CCRs and KCRs. Anion-selective ChRs (ACRs) conduct halides and nitrate, hyperpolarize the membrane in mature neurons, and inhibit their spiking (7). Cation-selective ChRs (CCRs) conduct primarily protons and, to a lesser extent, mono- and divalent metal cations (8). The relative permeability of CCRs for Na^+^ is greater than that for K^+^, so under physiological conditions they depolarize the membrane and activate neuronal spiking (9). Recently, we reported two “kalium channelrhodopsins” from the stramenopile fungus-like protist *Hyphochytrium catenoides* (*Hc*KCR1 and *Hc*KCR2) that are more permeable for K^+^ than Na^+^, and demonstrated that these light-gated channels can be used to inhibit mouse cortical neurons (10). Notably, KCRs lack the “K^+^ channel signature sequence” universally found in K^+^ channels from bacteria, archaea, eukaryotic cells and their viruses gated by voltage, ligands, heat, pH, or membrane deformation (11, 12). The ability of KCRs to discriminate between K^+^ and Na^+^ is particularly intriguing, because it reveals the only so far known alternative mechanism of K^+^ selectivity.

Cation conductance has appeared at least twice in microbial rhodopsin evolution, as CCRs from chlorophytes and streptophytes show very little protein sequence homology to CCRs from cryptophytes. Structurally and functionally the latter resemble haloarchaeal proton-pumping rhodopsins such as bacteriorhodopsin, and are therefore known as “bacteriorhodopsin-like cation channelrhodopsins”, or BCCRs (13). KCR protein sequences show the highest homology to cryptophyte BCCRs out of all currently known ChRs (10), although their source organism is phylogenetically very distant from cryptophytes. High-resolution structures of only one BCCR, known as ChRmine, have been reported (14, 15). They show trimeric organization typical of haloarchaeal ion-pumping rhodopsins (16), whereas chlorophyte CCRs and cryptophyte ACRs form dimers (17, 18).

In addition to two KCRs, the completely sequenced genome of *H. catenoides* encodes a third paralog (19). Surprisingly, this channel, named *H. catenoides* cation channelrhodopsin (*Hc*CCR), did not show higher permeability for K^+^ than for Na^+^ when tested by planar automated patch clamp (20). Thus, the three *H. catenoides* ChRs form a unique highly homologous group of light-gated channels with the K^+^/Na^+^ permeability ratios differing over a wide range. Here we used them as a platform to elucidate the structural foundations of the K^+^ selectivity mechanism of KCRs. In addition, we tested 13 homologs from other protists and metagenomic samples, and characterized those that are electrogenic. The results obtained confirmed our conclusions about the K^+^ selectivity mechanism drawn from analysis of *H. catenoides* ChRs.

## RESULTS

### Characterization of *Hc*CCR

Previously, we tested *Hc*CCR expressed in HEK293 (human embryonic kidney) cells by automatic patch clamp using complex solutions that do not allow a straightforward estimation of the K^+^/Na^+^ permeability ratio (P_K_/P_Na_) (20). Here we measured its current-voltage relationship (IV curve) by manual patch clamp under bi-ionic conditions (130 mM NaCl in the bath and 130 mM KCl in the pipette; for full solution compositions see Table S1), which we had used earlier to characterize *Hc*KCRs (10). Figure 1A shows a series of photocurrents generated by *Hc*CCR under incremental voltage, and Fig. 1C (red), the mean voltage dependence of the peak photocurrent. The reversal potential (V_rev_), estimated by approximation of the IV curve to zero current, was >40 mV under these conditions (Fig. 1C, red). Figure 1B shows a series of photocurrents generated by *Hc*CCR upon replacement of Na^+^ in the bath with K^+^. The mean IV curve measured with symmetrical K^+^ is shown in Fig. 1C (blue). The current amplitude was reduced, and the V_rev_ shifted to zero (Fig. 1C, blue). The P_K_/P_Na_ value, calculated from the V_rev_ shift using the Goldman-Hodgkin-Katz equation (21), was ∼0.2. This value was 128-fold and 94-fold smaller than P_K_/P_Na_ of *Hc*KCR1 and *Hc*KCR2, respectively (10), and even smaller than that of ChR2 from *C. reinhardtii*, the best characterized chlorophyte CCR (0.3-0.5 (8, 22)). Under bi-ionic conditions, *Hc*KCR1 showed a shift of V_rev_ to more depolarized values during 1-s illumination (10). This shift was even larger in the recently identified KCR from the stramenopile *Wobblia lunata*, named *W*iChR1 for “*Wobblia* inhibitory ChR1” (Fig. 1D, black; Fig. S1; and (23)). However, no such shift was detected in *Hc*CCR (Fig. 1D, red), which confirmed our previous conclusion that in *Hc*KCR1 this shift reflected a decrease in P_K_/P_Na_ during illumination (10). The decay of *Hc*CCR photocurrent slightly accelerated upon depolarization, but was slower than that in both *Hc*KCRs (Fig. 1E). The maximal spectral sensitivity of *Hc*CCR was at 530 nm (Fig. 1F).

**FIG 1.**
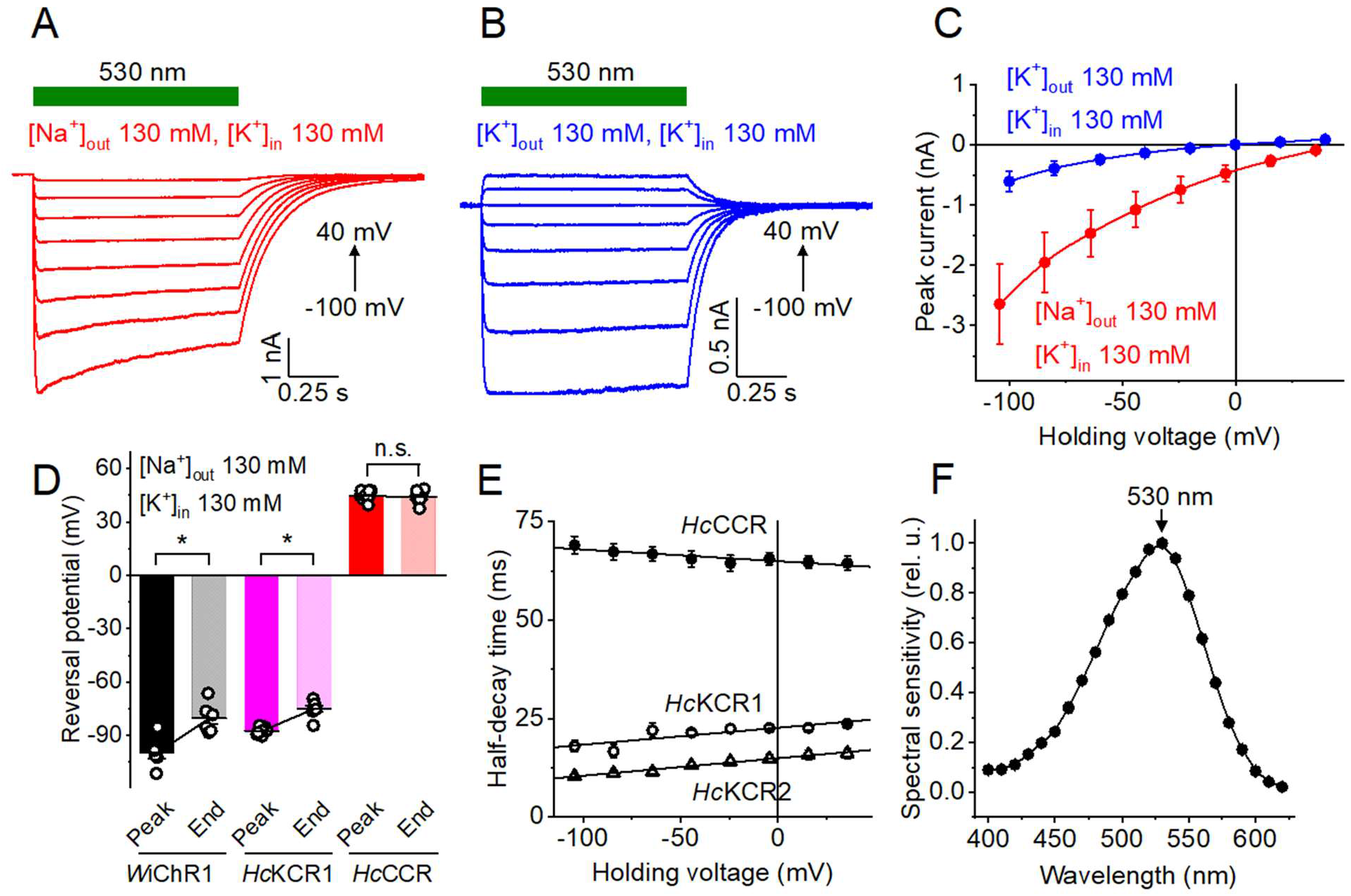
Electrophysiological characterization of *Hc*CCR. (A and B) Series of photocurrent traces recorded from *Hc*CCR upon incremental voltage with 130 mM Na^+^ (A) or K^+^ (B) in the bath and 130 mM K^+^ in the pipette. The duration of the light pulse is shown by the green bars. (C) The peak current-voltage relationships of *Hc*CCR under the indicated ionic conditions. The data points are the mean ± sem (n = 7 and 6 cells, for the Na^+^ and K^+^ bath, respectively). (D) The reversal potentials of the peak current and current at the end of 1-s illumination measured in the Na^+^ bath as in A. The data points are the mean ± sem (n = 6-8 cells for each variant). *, p < 0.05 by the two-tailed paired sample Wilcoxon signed rank test; n.s., not significant. The data for *Hc*KCR1 and *Hc*KCR2 are shown for comparison. (E) The dependence of the photocurrent half-decay time on the holding voltage for the three *H. catenoides* ChRs. The data points are the mean ± sem (n = 6 cells for each variant), the lines are linear approximations. (F) The action spectrum of *Hc*CCR photocurrents. The data points are the mean ± sem (n = 6 cells). The numerical data for panels C, D, E and F, including the exact numbers of cells sampled, are provided in Data Set 1, and full statistical analysis including the exact p values, in Data Set 4.

### *Hc*CCR_*Hc*KCR1 chimeras and mutants

The seven-transmembrane (7TM) domain of *Hc*CCR shares 70-73% identity and 83-86% similarity at the protein level with those of KCRs (Fig. S2). Remarkably, the protein alignment shows no gaps, so that the numbers of the homologous residues are the same in all three proteins. As the first step towards determination of the structural foundations of the K^+^ selectivity of *Hc*KCRs, we carried out patch clamp analysis of *Hc*CCR_*Hc*KCR1 chimeras. Starting with the *Hc*CCR sequence, we systematically replaced individual predicted helical regions with those of *Hc*KCR1. We have also created an additional chimera by replacement of the N-terminal region of *Hc*CCR with that of *Hc*KCR1. A protein alignment of the chimeras is shown in Fig. S3, and their schematic representation, in Fig. 2A. Next, we measured the IV curves of the chimeras under the bi-ionic conditions (Fig. S4) and calculated the V_rev_ values as described above for wild-type *Hc*CCR. Remarkably, replacement of TM2 or TM7 caused a >40-mV shift of V_rev_ to more negative values, indicating a large increase in P_K_/P_Na_ (Fig. 2B). These results suggested that residues responsible for the K^+^ selectivity of *Hc*KCRs are located in TM2 and TM7.

**FIG 2.**
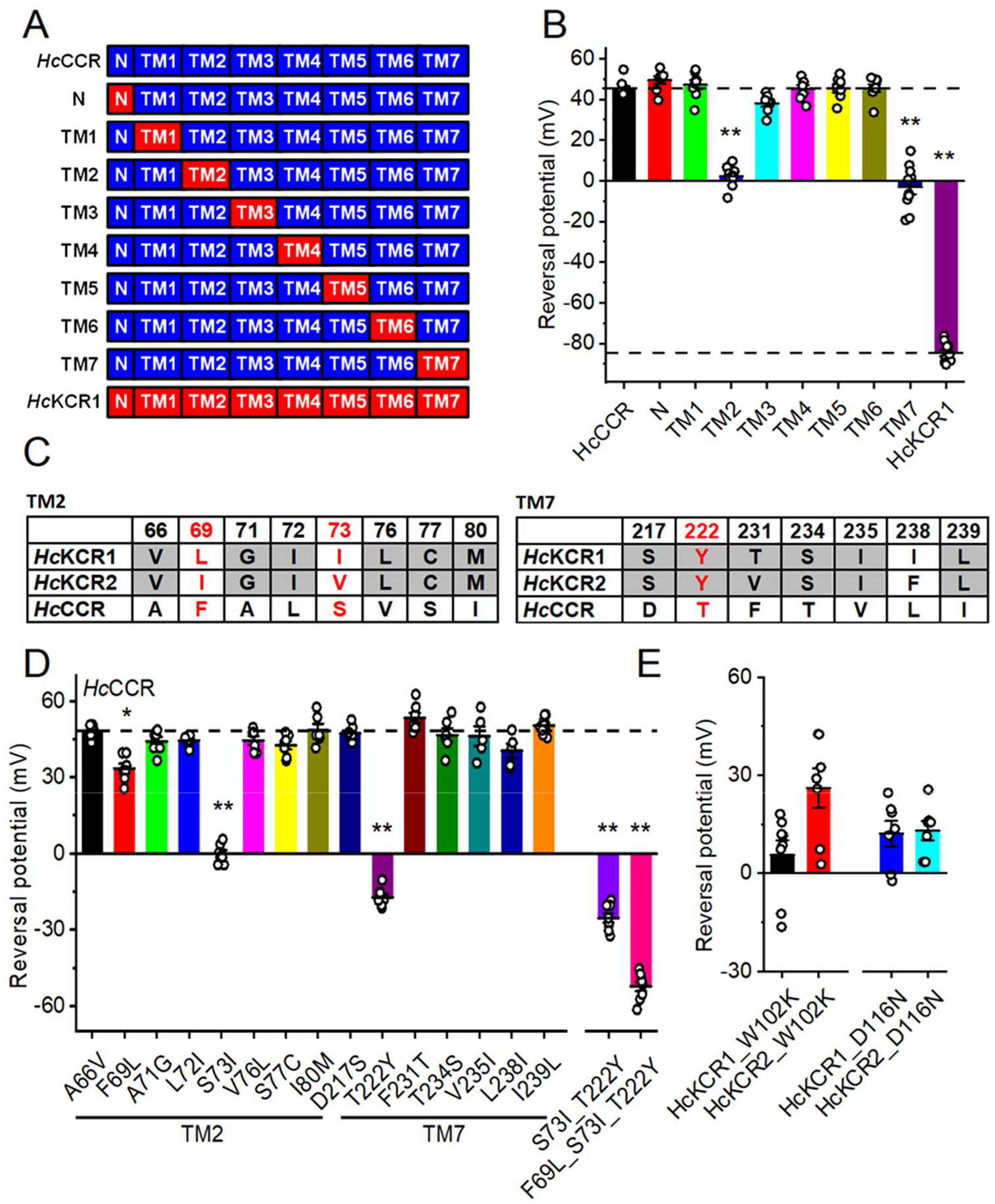
Analysis of *Hc*CCR chimeras and mutants. (A) A schematic representation of *Hc*KCR1_*Hc*CCR chimeras tested in this study. (B) The V_rev_ values measured in the wild-type proteins and chimeras under bi-ionic conditions (130 mM Na^+^ in the bath and 130 mM K^+^ in the pipette). The dashed lines mark the V_rev_ value of wild-type *Hc*CCR and *Hc*KCR1. (C) The residues in the tested positions in TM2 and TM7 of the three *H. catenoides* ChRs. The red font shows the positions critical for K^+^ selectivity. (D) The V_rev_ values measured in the single and multiple *Hc*CCR mutants as described in B. The dashed line marks the V_rev_ value of wild-type *Hc*CCR. (E) The V_rev_ values measured in the W102K and D116N mutants of *Hc*KCR1 and *Hc*KCR2. In B, D and E, the bars and whiskers show the mean ± sem (n = 5-10 cells); the empty circles, the data for individual cells. *, p < 0.05, **, p < 0.01 by one-way ANOVA followed by the Tukey test for means comparison. The numerical data for panels B, D and E are provided in Data Set 2, and full statistical analysis, in Data Set 4. and full statistical analysis including the exact p values, in Data Set 4.

Next, we identified the positions in TM2 and TM7 occupied by the same residue in both *Hc*KCRs but not *Hc*CCR, and the positions in which the residues are different in all three proteins (Fig. 2C). We individually replaced the residues of *Hc*CCR with those found in *Hc*KCR1, and measured the IV curves of the resultant point mutants under the bi-ionic conditions (Fig. S5). Only three mutations (F69L, S73I, and T222Y) caused a significant shift of V_rev_ towards more negative values (Fig. 2D), indicating that the mutated residue positions are critical for the K^+^ selectivity of *Hc*KCRs. The effect of the individual mutations was synergistic, as the single F69L mutation caused a larger V_rev_ when it was added to the *Hc*CCR_S73I_T222Y double mutant than when it was made in wild-type *Hc*CCR (Fig. 2D). The three mutations together (F69L, S73I and T222Y) converted Na^+^-selective *Hc*CCR into a KCR with P_K_/P_Na_ ∼8. In *Hc*KCR2, which shows a slightly lower P_K_/P_Na_ than *Hc*KCR1 (10), the position 73 is occupied by Val instead of Ile. The *Hc*KCR2_V73I mutation caused a small, but statistically significant shift of the V_rev_ from -74 ± 2 to -79 ± 1 mV (mean ± sem, n = 7 and 8 cells for the wild type and the mutant, respectively, p = < 0.05 by the two-tailed Mann-Whitney test; for full statistical analysis see Data Set 4), which corresponded to an increase of P_K_/P_Na_ from 17 to 22.

Arginine in the position 82 of bacteriorhodopsin contributes to its complex counterion to the protonated retinylidene Schiff base (24) and is highly conserved in all microbial rhodopsins, including chlorophyte and streptophyte CCRs. As an exception, in BCCRs the prevalent residue in the corresponding position is Lys, and in all three *H. catenoides* paralogs the corresponding position is occupied with Trp102. The *Hc*KCR1_W102R mutation completely abolished photocurrents, whereas *Hc*KCR1_W102K did generate small currents (Fig. S6). Surprisingly, this mutant showed a small positive V_rev_ under our bi-ionic conditions (Fig. 2E, black), reflecting a dramatic decrease in P_K_/P_Na_ caused by the mutation. An even more positive V_rev_ was observed in the *Hc*KCR2_W102K mutant (Fig. 2E, red).

The Asp residue in the Schiff base proton donor position (corresponding to Asp96 in bacteriorhodopsin) is conserved in all three *H. catenoides* paralogs, as in most cryptophyte BCCRs. Mutagenetic neutralization of this residue strongly inhibited photocurrents in both *Hc*KCRs (Fig. S6), and shifted the V_rev_ to more positive values (Fig. 2E, blue and cyan), indicating a decrease in P_K_/P_Na_. We conclude that Trp102 and Asp116 are necessary, although not sufficient, for the K^+^ selectivity of *Hc*KCRs. The Asp116 mutations, but not the W102 mutations, also caused an inward rectification of the IV curves in both *Hc*KCRs (Fig. S6).

### Homology modeling of *Hc*KCR1 and *Hc*CCR

To gain an insight into locations of the critical residues identified in the previous section and predict their possible interactions, we created homology models of *Hc*KCR1 and *Hc*CCR (Fig. 3). The root-mean-square deviation (RMSD) of atomic positions between the two models is 0.7 Å. In both models, the residues 69 and 73 are located in the cytoplasmic half of TM2 in the vicinity of Asp116, and the residue 222, near the extracellular surface of the protein within 5 Å of Trp102. All these residues are expected to contribute to the putative cation conduction pathway formed by TM1, 2, 3 and 7, as in other ChRs. In the *Hc*CCR model, the orientation of the Trp102 sidechain is rotated upward from that in *Hc*KCR, likely as the result of the substitution of a more compact Thr for Tyr in the position 222. This conformational difference, if confirmed by X-ray crystallography or cryo-electron microscopy (cryo-EM), may be relevant for control of P_K_/P_Na_. Empirical calculations (25) predict that at pH 7.4 Asp116 is unprotonated in both channels (pK_a_ ∼4). In both models, Asp116 forms sidechain hydrogen bonds with Ser70 in the middle of TM2 and Arg 244 at the cytoplasmic end of TM7. In *Hc*KCR1, the S70A mutation decreased P_K_/P_Na_ (23), but Ser70 is conserved in *Hc*CCR and does not render this channel K^+^-selective. Considering the results of our mutant analysis (Fig. 2D), it is plausible that properties of Ser70 in *Hc*CCR compared to those in *Hc*KCR1 are modified by the substitution of Phe for Leu, and Ser for Ile in the nearby positions 69 and 73, respectively. The latter, polar-to-nonpolar substitution, produced a particularly large effect on the channel selectivity.

**FIG 3.**
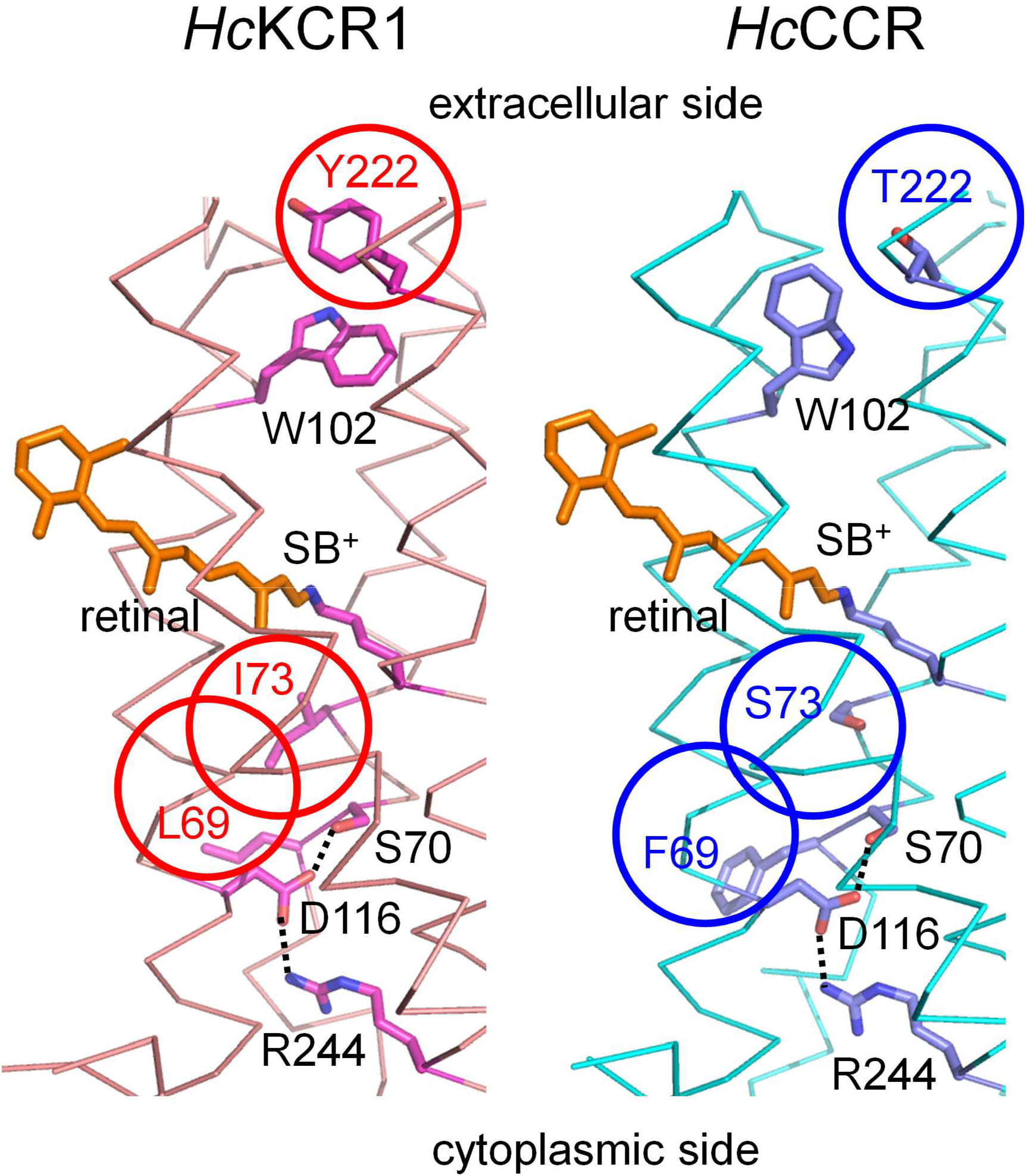
Homology models of *Hc*KCR1 (left) and *Hc*CCR (right). The sidechains of the critical residues and the retinal chromophore are shown as sticks; the transmembrane helices, as ribbons; the predicted hydrogen bonds, as dotted lines. SB^+^, protonated Schiff base. The three residues responsible to the difference in the relative permeability are circled.

### KCR orthologs from other sources

To verify our conclusions about the structural foundations of the K^+^ selectivity drawn from the analysis of *Hc*CCR_*Hc*KCR1 chimeras and mutants, we searched the genomes and transcriptomes of other microorganisms and environmental samples for orthologs of *H. catenoides* ChRs. We identified 13 sequences that encode rhodopsin domains clustering together with *H. catenoides* ChRs on a phylogenetic tree (Fig. 4A). Among these, two sequences were found in the predatory alveolate *Colponema vietnamica* (26); one sequence, in the bicosoecid strain BVI, formerly considered as *Cafeteria roenbergensis* but recently reattributed as *C. burkhardae* (Dr. Matthias Fischer, Max Planck Institute for Medical Research, personal communication); three sequences, in *Chromera velia*, an alga related to apicomplexan parasites (27); and seven sequences, in various metagenomic databases (listed in Methods). A protein alignment of their rhodopsin (7TM) domains is shown in Fig. S7. The three *C. velia* sequences, and metagenomic MATOU-v2.32008995_3 and TARA_MED_95_MAG_00407_000000002956 differ from the rest by a non-carboxylate residue in the counterion position (corresponding to Asp85 in bacteriorhodopsin). A very unusual feature of the three *C. velia* sequences is substitution of Gly for Thr89 (bacteriorhodopsin numbering) that is nearly universally replaced with Cys in other known ChRs.

**FIG 4.**
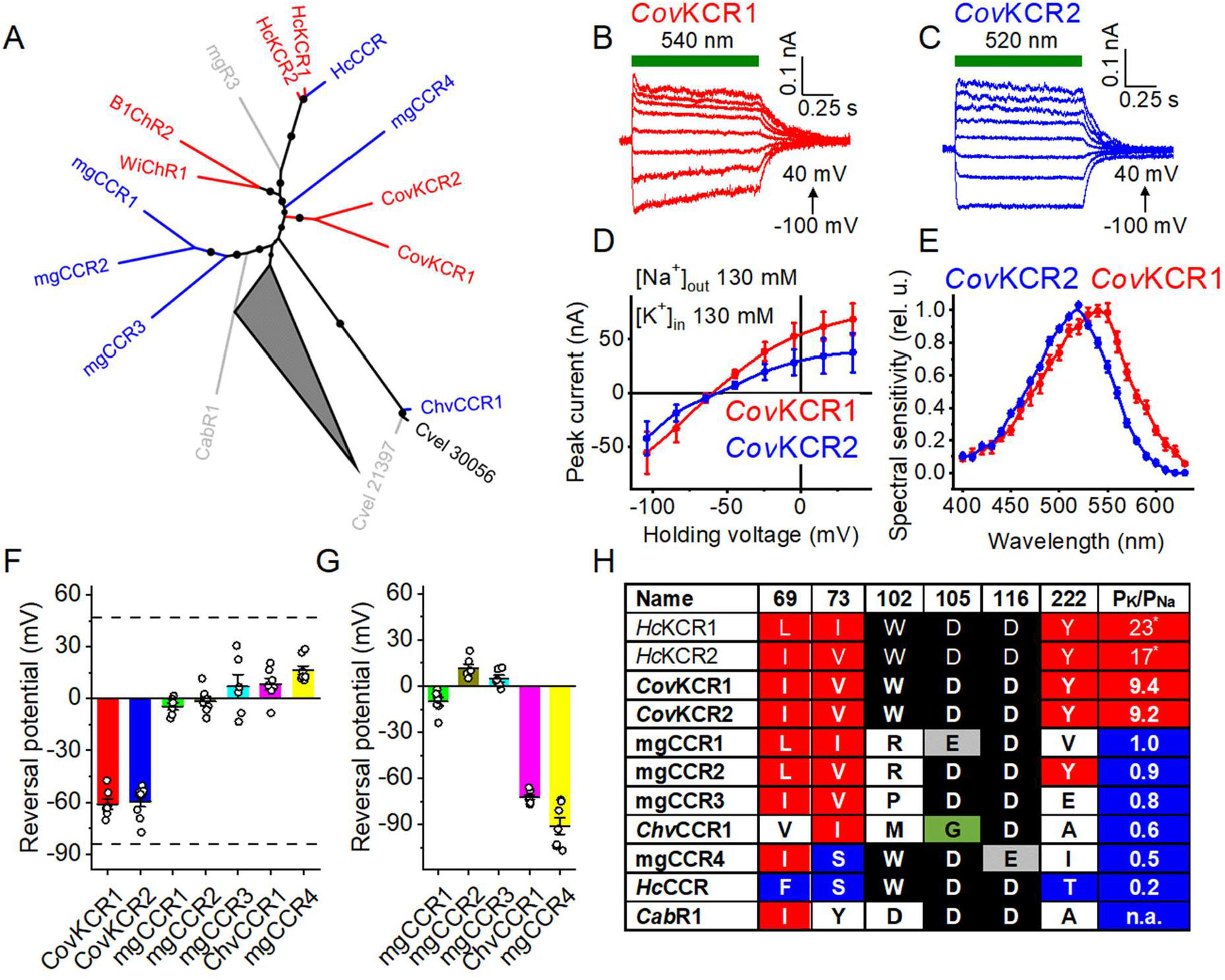
Characterization of KCR homologs. (A) A phylogenetic tree of the tested *Hc*KCR homologs. Variants that exhibit the K^+^ selectivity are in red, those that do not, in blue, and variants that do not generate channel currents, in grey. The collapsed node shows 510 other known ChRs (including non-functional mgR1 and mgR2). The 50-100 bootstrap values are shown as circles. (B and C) Series of photocurrent traces recorded from *Cov*KCRs upon incremental voltage with 130 mM Na^+^ in the bath and 130 mM K^+^ in the pipette. The green bars show the duration of the light pulse. (D) The IV curves of *Cov*KCRs measured under these conditions. The data points are mean ± sem, n = 7 and 9 cells for *Cov*KCR1 and *Cov*KCR2, respectively. (E) The action spectra of *Cov*KCR photocurrents. The data points are mean ± sem, n = 7 cells for each variant. (F and G) The V_rev_ values of all tested homologs measured under these conditions (F) and upon replacement of Na^+^ in the bath with NMDG^+^. The bars and whiskers show the mean ± sem (n = 6-9 cells); the empty circles, the data for individual cells. (H) The residues in the positions important for the K^+^ selectivity. Bold text shows variants tested in this study. *, the values are from (10); n.a., not applicable. The red and blue backgrounds highlight the residues that, respectively, increase or decrease the K^+^ selectivity. The green background shows the non-carboxylate residue in the proton acceptor position in the *C. velia* sequence. The numerical data for panels D, E, F and G, including the exact numbers of cells sampled for each variant, are provided in Data Set 3.

We synthesized mammalian codon-adapted versions of polynucleotides encoding these rhodopsins and expressed them as mCherry fusions in human embryonic kidney (HEK293) cells. Four homologs (Cvel_21397, MATOU-v2.141421879_4, MATOU-v2.32008995_3, and TARA_MED_95_MAG_00407_000000002956) were very poorly expressed and generated no photocurrents. We assigned the names mgR1-mgR3 (metagenomic rhodopsins 1-3) to the sequences MATOU-v2.141421879_4, MATOU-v2.32008995_3, and TARA_MED_95_MAG_00407_000000002956, respectively. Cvel_30056 generated photocurrents, but they were smaller than those from the closely related Cvel_28437, so we did not characterize them. Photocurrents from Cvel_28437 (which we named *Chv*CCR1) and all other homologs except KAA0157615 from *C. burkhardae* demonstrated voltage-dependent sign reversal characteristic of passive conductance. We measured the IV curves under bi-ionic conditions, as described above for *Hc*CCR. Only two homologs (GILI01001652 and GILI01010992 from *C. vietnamica*) showed negative V_rev_ values, indicating their higher permeability for K^+^ over Na^+^. Fig. 4B and 4C show series of their photocurrent traces, and Fig. 4D, the corresponding IV curves. The action spectrum of GILI01001652 photocurrents peaked near 540 nm, and that of GILI01010992, at 520 nm (Fig. 4E). Following the general ChR numbering convention, we named the more red-shifted paralog *Cov*KCR1, and the more blue-shifted one, *Cov*KCR2. Representative photocurrent traces, the corresponding IV curves and the spectra of other functional homologs are shown in Fig. S8. All functional homologs showed inward rectification, also typical of many chlorophyte CCRs and cryptophyte BCCRs. We assigned the names mgCCR1-mgCCR4 to the sequences Ga0170791_133102851, Ga0007756_110676931, MATOU-v2.119411731_5 and Ga0392354_009429_356_1807, respectively. When Na^+^ in the bath was replaced with non-permeable *N*-methyl-*D*-gluconate (NMDG^+^), the IV curves for *Chv*CCR1 and mgCCR4 showed large shifts to the left, indicating a substantial permeability for Na^+^ (Fig. S9). The IV curves for mgCCR1-mgCCR3 showed little change, indicating that these homologs primarily conduct other ions, most probably, H^+^. When the pH of the bath was raised to 9.4, the IV curves of all these three channels shifted to more negative voltages (Fig. S10), which confirmed this hypothesis. The V_rev_ values measured in the Na^+^ and NMDG^+^ baths are shown in Fig. 4F and G, respectively.

Fig. 4H shows the residues in the critical positions in the sequences tested in this study and the earlier characterized *Hc*KCRs. Only the K^+^-selective *C. vietnamica* sequences contain all residues identified as important for K^+^ selectivity in *Hc*KCRs, whereas in the non-K^+^-selective homologs some corresponding positions are occupied with non-homologous residues. This strongly supports our conclusions about the structural determinants of K^+^ selectivity in ChRs. In particular, our results show that the presence of both the Trp102 homolog and the Tyr222 homolog is required for discrimination between K^+^ and Na^+^. Neither mgCCR4, in which only Trp102 is conserved but Tyr222 is replaced with Ile, nor mgCCR2, in which only Tyr222 is conserved but Trp102 is replaced with Arg, exhibit K^+^ selectivity.

Positive photocurrents from KAA0157615 recorded in the Na^+^ bath decayed with a time constant of ∼15 ms after the onset of illumination and were practically independent of voltage (Fig. S8). This behavior is typical of active intramolecular proton transfer from the Schiff base to an outwardly located acceptor (28). We assigned the name *Cab*R1 (*Cafeteria burkhardae* rhodopsin 1) to this protein.

## DISCUSSION

ChRs are found in many taxa of eukaryotic microbes, both photosynthetic and heterotrophic (1). During the last 17 years, ChRs served as extremely powerful and versatile tools for optical control of the membrane potential in excitable cells such as neurons and myocytes, and thus have become indispensable for neuroscience research (29). Moreover, partial recovery of visual function in a blind human patient by optogenetic means has launched a new era of ChR gene therapy (30). Our recent discovery of natural ChRs with a high P_K_/P_Na_ ratio (KCRs) (10) complements the inventory of optogenetic tools with long-sought, nearly universal inhibitory molecules.

Despite their prominence in biomedicine, molecular mechanisms of ChRs, and especially the structural foundations of their ionic selectivity, are still poorly understood. Unlike conventional ion channels gated by voltage or ligands, in which the ion conductance pathway is formed at the interface between several subunits, it appears that each individual ChR protomer is capable of ion conductance. An inter-protomer conductance has been proposed in ChRmine, based on the cryo-EM structure obtained in detergent (14). However, lipids block the space between the protomers in a ChRmine trimer incorporated in membranous nanodisks (15), which is expected to represent a state of the protein close to that in biological membranes.

In this study, we have taken advantage of the existence of closely homologous ChRs, *Hc*KCR1 and *Hc*CCR, that differ >100-fold in their P_K_/P_Na_. By systematic replacement of individual transmembrane helices of *Hc*CCR with those of *Hc*KCR1, we found that TM2 and TM7 are responsible for the K^+^ selectivity. These helices contribute to the formation of the ion conduction pathway and channel gating in other ChRs, as shown by electron paramagnetic resonance (31, 32) electron crystallography (33), and X-ray crystallography (34). Then, we systematically mutated all divergent residues in TM2 and TM7 of *Hc*CCR, replacing them with those found in *Hc*KCR1 in the corresponding positions. We identified three residue positions, two in TM2 (69 and 73) and one in TM7 (222), critical for the K^+^ selectivity of the *Hc*KCRs. A lower P_K_/P_Na_ of the triple mutant *Hc*CCR_F69L_S73I_T222Y than that of the wild-type *Hc*KCR1 is likely explained by synergy of other divergent residues that produce no significant effect individually, but increase P_K_/P_Na_ in combination with other mutations, as we found for F69L. Phe/Leu69 and Ser/Ile73 are the homologs of Val49 and Ala53 of bacteriorhodopsin, respectively. In the latter protein, these residues control the position of the Schiff base lysine (Lys216) sidechain relative to Asp85 (conserved in *H. catenoides* ChRs as Asp105) and affect distribution of the proton between them (35). In chlorophyte CCRs, the position of Ser/Ile73 is occupied by a highly conserved Glu residue (Glu90 in *Cr*ChR2). It contributes to the “central gate” and controls the selectivity of the channel: mutation of Glu90 to Lys or Arg renders *Cr*ChR2 permeable to anions (36). Considering the low protein sequence homology between *H. catenoides* ChRs and chlorophyte CCRs, the importance of this position in determination of the channel selectivity suggests a functional principle common to all ChRs.

Remarkably, two residues that are conserved in all three *H. catenoides* ChRs, namely, Trp102 and Asp116, are also required for the K^+^ selectivity of *Hc*KCRs, as we found by testing their W102K and D116N mutants. Trp102 corresponds to Arg82 of bacteriorhodopsin, which is highly conserved in microbial rhodopsins. Mutagenetic neutralization of the homologous residue (the R109N mutation) in the Na^+^-pumping rhodopsin from the flavobacterium *Dokdonia eikasta* (KR2) brings about weak passive K^+^ conductance (37). In cryptophyte BCCRs, the Arg82 position can be occupied by Pro as in *Gt*CCR1 and *Gt*CCR2 from *Guillardia theta* (13), or even Glu, as in *Ra*CCR2 from *Rhodomonas abbreviata* (38), but not by Trp, as in KCRs.

Our homology model shows a close proximity of Trp102 and Tyr222 positions in the extracellular portion of the putative cation pathway within *Hc*KCR1. In *W*iChR1 and a KCR from the stramenopile *Bilabrum* sp. (*B1*ChR2; (23)), the residue corresponding to Tyr222 of *Hc*KCRs is Phe. Both these KCRs showed higher P_K_/P_Na_ values than *Hc*KCR1, but the *Hc*KCR1_Y222F mutant exhibited a decrease rather than increase in the K^+^ selectivity (23). Therefore, the Phe for Tyr substitution is unlikely responsible for the larger P_K_/P_Na_ of *W*iChR1 and *B1*ChR2, as compared to that of *Hc*KCR1. Several conserved aromatic residues are also found in the pore region of animal voltage-gated K^+^ channels, and the cation-π interaction has been proposed to contribute to their selectivity (39). A similar mechanism can be at work in microbial KCRs.

Asp116 of *Hc*KCRs corresponds to Asp96 of bacteriorhodopsin, the proton donor during reprotonation of the Schiff base (40). In chlorophyte CCRs, this Asp is replaced with a non-carboxylate residue (His173/His134 in *Cr*ChR1/*Cr*ChR2), and the *Cr*ChR1_H173D mutation completely abolished channel currents (41). In cryptophyte BCCRs, this Asp is conserved as in KCRs. In *Gt*CCR2 (one of BCCRs), deprotonation of the Asp96 homolog (Asp98) occurs >10-fold faster than reprotonation of the Schiff base and is required for cation channel opening (13). When we were preparing our manuscript for submission, a preprint was published reporting patch clamp analysis of *Hc*KCR1 mutants (23). Its results provide an independent validation of the conclusions drawn in our study. In addition to Ser70, Trp102 and Asp116, described above, mutations of Asp87 and Asn99 also reduced P_K_/P_Na_ in *Hc*KCR1 (23), although both these residues are conserved in Na^+^-selective *Hc*CCR. High-resolution structures are likely needed to explain this observation.

The P_K_/P_Na_ ratios of mammalian voltage-gated K^+^ channels fall within the range 100-1,000 (12), which is higher than that of microbial KCRs. Our identification of the residues required for the K^+^ selectivity of ChRs is expected to facilitate both bioinformatic searches for potentially highly K^+^-selective ChR sequences and their molecular engineering to further improve the K^+^ selectivity. *Cov*KCR1 and *Cov*KCR2 tested in this study are relatively poor candidates for the development of optogenetic tools, as their P_K_/P_Na_ are lower than those of *Hc*KCRs, and their photocurrents are very small. Nevertheless, these KCRs confirm the importance of the presence of both Trp102 and Tyr222 homologs for the K^+^ selectivity. Also, their source organism, *C. vietnamica*, is phylogenetically very distant from *H. catenoides*, which suggests a wide distribution of KCRs in eukaryotic taxa. On the other hand, species attribution of transcripts derived from predatory microorganisms, such as *Colponema*, should be treated with caution, as there is a possibility of contamination of their transcriptomes with RNA from food. This happened, e.g., when ChRmine, the sequence encoded by the *Cryptomonas lens* genome, was erroneously attributed to the ciliate *Tiarina fusus* fed on *C. lens* (as discussed in (38)).

Intramolecular proton transfers preceding channel currents were detected earlier in some chlorophyte CCRs (28) and cryptophyte BCCRs (13). Several chlorophyte sequences highly homologous to ChRs generates only fast photocurrents reflecting these transfers but shows no passive ion conductance upon expression in mammalian cells (1), similar to *Cab*R1 described here. One possible explanation of the lack of channel activity in these proteins observed in heterologous systems is that they are more sensitive to membrane components, e.g., the lipid composition of the membrane, than other ChRs.

The genome of *Cafeteria* strain BVI (as well as those of strains E4-10P and Cflag) also encodes a functional ACR that shows closer homology to haptophyte ACRs than to ACRs from other stramenopiles (1), suggesting horizontal gene transfer. The presence of rhodopsins that belong to different families and functional types is common in many taxa, including chlorophytes, cryptophytes and dinoflagellates. The genome of *C. velia* also encodes several other rhodopsin sequences besides the three tested in this study, most of which cluster together with non-electrogenic rhodopsins from other organisms, and their functional roles remain unknown. Only a small fraction of protist rhodopsins have been characterized at any level beyond their primary structure, and further characterization will likely bring many striking discoveries. A recent example are algal rhodopsins that act as photosensitive domains to regulate a pentameric bestrophin channel (42).

## MATERIALS AND METHODS

### Bioinformatics and molecular biology

*Hc*KCR homologs were identified by BLAST (BLASTP and TBLASTN) searches of various public databases, using a truncated TM domain amino acid sequence of *Hc*KCR1 (residues 13-251) as a query. Specifically, GILI01001652 and GILI01010992 proteins of *C. vietnamica* strain Colp-7a were found using NCBI TBLASTN against the Transcriptome Shotgun Assembly (TSA) database limited to the SAR supergroup (taxid:2698737). KAA0157615 protein of *Cafeteria burkhardae* strain BVI was found using NCBI BLASTP against the Non-redundant (NR) protein database. Metagenomic MATOU-v2.141421879_4, MATOU-v2.32008995_3, MATOU-v2.119411731_5, and TARA_MED_95_MAG_00407_000000002956 proteins were found using BLASTP in the Ocean Gene Atlas (https://tara-oceans.mio.osupytheas.fr/ocean-gene-atlas/) (43, 44). The MATOU-v2.141421879_4, MATOU-v2.32008995_3, and MATOU-v2.119411731_5 were extracted from the Marine Atlas of Tara Ocean Unigenes (MATOU; (45)) dataset, while TARA_MED_95_MAG_00407_000000002956 was found in Tara Oceans Single-Cell and Metagenome Assembled Genomes (EUK-SMAGs;(46)) dataset. Metagenomic Ga0392354_009429_356_1807, Ga0007756_110676931, and Ga0170791_133102851 proteins were found in the Department of Energy (DOE) Joint Genome Institute (JGI) Integrated Microbial Genomes and Microbiomes (IMG/M) database (47). BLASTP search was used against the respective metatranscriptomes (Metatranscriptome of lab enriched marine microbial communities from Marineland, Florida, USA - SWA_R2_TP1; Metatranscriptome of freshwater lake microbial communities from Lake Michigan, USA - Su13.BD.MLB.DD; and Northern Canada Lakes metatranscriptome co-assembly). Finally, amino acid sequences for *Chromera velia* proteins Cvel28437, Cvel21397, Cvel30056 were found in PhycoCosm (https://phycocosm.jgi.doe.gov/Chrveli1/Chrveli1.home.html) (48) using BLASTP against Chromera_velia 20200809 filtered protein models dataset (49). Polynucleotides encoding the 7TM domains of the predicted proteins were optimized for mammalian expression. For expression in HEK293 (human embryonic kidney) cells these polynucleotides and those encoding *H. catenoides* ChRs (GenBank accession numbers MZ826861, MZ826862 and OL692497) were cloned into the mammalian expression vector pcDNA3.1 (Life Technologies) in frame with a C-terminal mCherry tag.

The transmembrane helices were predicted using the DeepTMHMM algorithm (50). Sequences were aligned using MegAlign Pro software v. 17.1.1 (DNASTAR Lasergene) with default parameters. Phylogeny was analyzed with IQ-TREE v. 2.1.244 using automatic model selection and ultrafast bootstrap approximation (1000 replicates) (51). iTOL v. 6.346 was used to visualize and annotate phylogenetic trees.

### Homology modeling of *Hc*KCR1 and *Hc*CCR

We used ColabFold (52) with standard settings to generate, for each protein, five structural models based on multiple sequence alignments. The pLDDT (predicted Local Distance Difference Test) confidence scores (52) of all models were in a relatively narrow range, 80.8-82.0 for *Hc*KCR1, and 81.8-83.4 for *Hc*CCR. A single structural model was chosen for each protein based on the overall similarity of the scores and the results of manual inspection for details of secondary structure and local interactions. We used 5awz (the crystal structure of *Acetabularia* rhodopsin 1 (53)) selected by ColabFold for model building to dock the retinal chromophore and internal water molecules to the ColabFold models by aligning them with 5awz in PyMol. We kept seven internal water molecules for *Hc*KCR1, and six for *Hc*CCR. Coordinates for missing hydrogen atoms were generated with CHARMM (Chemistry at HARvard Molecular Mechanics) (54). The retinal-bound models of *Hc*KCR1 and *Hc*CCR with internal water molecules were subjected to geometry optimizations using the CHARMM potential energy function with CHARMM36 parameters for water (55, 56), TIP3P water model (57), and retinal parameters as described in (58-60). To optimize the geometry of retinal, water molecules, and protein groups within 3.5 Å of retinal and water, we fixed the coordinates of the heavy atoms of all other protein sidechains; on the heavy atoms of the mobile groups, we initially placed harmonic constraints of 10 kcal mol^-1^ Å^-2^ and energy-optimized to a gradient of 0.1 kcal mol^-1^ Å^-2^; we lowered the harmonic constraints first to 1.0 kcal mol^-1^ Å^-2^, then to 0.1 kcal mol^-1^ Å^-2^, each time performing a new energy optimization to a gradient of 0.1 kcal mol^-1^ Å^-2^. All harmonic constraints were then switched off and an additional energy optimization step applied.

### Whole-cell patch clamp recording from HEK293 cells

No cell lines from the list of known misidentified cell lines maintained by the International Cell Line Authentication Committee were used in this study. The HEK293 cells from the American Type Culture Collection (ATCC) were grown in 2-cm diameter plastic dishes and transfected with 10 μl of ScreenFectA transfection reagent (Waco Chemicals USA, Richmond) using 3 μg DNA per dish. Immediately after transfection, all-*trans*-retinal (Sigma) was added at the final concentration of 5 µM. Patch pipettes with resistances of 2-3 MΩ were fabricated from borosilicate glass. Whole-cell voltage clamp recordings were performed with an Axopatch 200B amplifier (Molecular Devices) using the solutions, full composition of which is shown in Table S1, and a 4 M salt bridge. All measurements were carried out at room temperature (25° C). The signals were digitized with a Digidata 1440A controlled by pClampEx 10.7 software (both from Molecular Devices). All current-voltage curves (IV dependencies) were corrected for liquid junction potentials (LJP) calculated using the ClampEx built-in LJP calculator (Table S1). A Polychrome IV light source (T.I.L.L. Photonics GMBH) in combination with a mechanical shutter (Uniblitz Model LS6, Vincent Associates; half-opening time 0.5 ms) was used as the light source (maximal quantum density at the focal plane of the 40× objective lens was ∼7 mW mm^-2^ at 540 nm). The action spectra were constructed using the initial slope of photocurrent in the linear range of the dependence on the quantum density (<25 µW mm^-2^), corrected for the quantum density measured at each wavelength and normalized to the maximal value. ClampFit 10.7 was used for Initial analysis of the recorded data, followed by further analysis by Origin Pro 2016 software (OriginLab Corporation). The data points shown in the graphs are connected with spline or B-spline lines, unless otherwise stated.

### Statistics and reproducibility

Identical batches of HEK293 cell culture were randomly assigned for transfection with each tested construct. At least two separate batches of culture were transfected independently with each construct. Individual transfected cells were selected for patching by inspecting their tag fluorescence. Non-fluorescent cells, or cells in which no GΩ seal could be established were not sampled. Only one photocurrent trace per cell was recorded, and traces recorded from different cells transfected with the same construct were considered biological replicates (reported as n values). Statistical analysis was performed using Origin Pro 2016 software. The data are presented as mean ± sem values, the data from individual cells are also shown when appropriate. Normal distribution of the data was checked using the Kolmogorov-Smirnov test. Specific statistical hypotheses were tested using the two-tailed paired sample Wilcoxon signed rank test, the two-tailed Mann-Whitney test, and one-way ANOVA followed by the Tukey test for means comparison, as implemented in Origin. The complete results of hypothesis testing (including the number of cells tested for each variant and the exact p values) are provided in Data Set 4.

### Data availability

The polynucleotide sequences of KCR homologs reported in this study have been deposited to GenBank (accession numbers OP121639-OP121651). The numerical values of the data shown in Figs. 1, 2 and 4 are provided in Data Sets 1-3, respectively.

## Acknowledgements

We thank Dr. Oded Béjà (Technion-Israel Institute of Technology, Haifa, Israel) for sharing with us the protein sequence information of *W*iChR1 and *B1*ChR2 prior to peer-reviewed publication, and Dr. Matthias Fischer (Max Planck Institute for Medical Research, Heidelberg, Germany) for the species attribution of the KAA0157615 sequence, and Dr. Oliver P. Ernst (University of Toronto, Canada) for critical reading of the manuscript. We also thank Dr. Hai Li from the Spudich lab for helpful discussions and Yumei Wang for technical assistance.

---

**Table S1.**
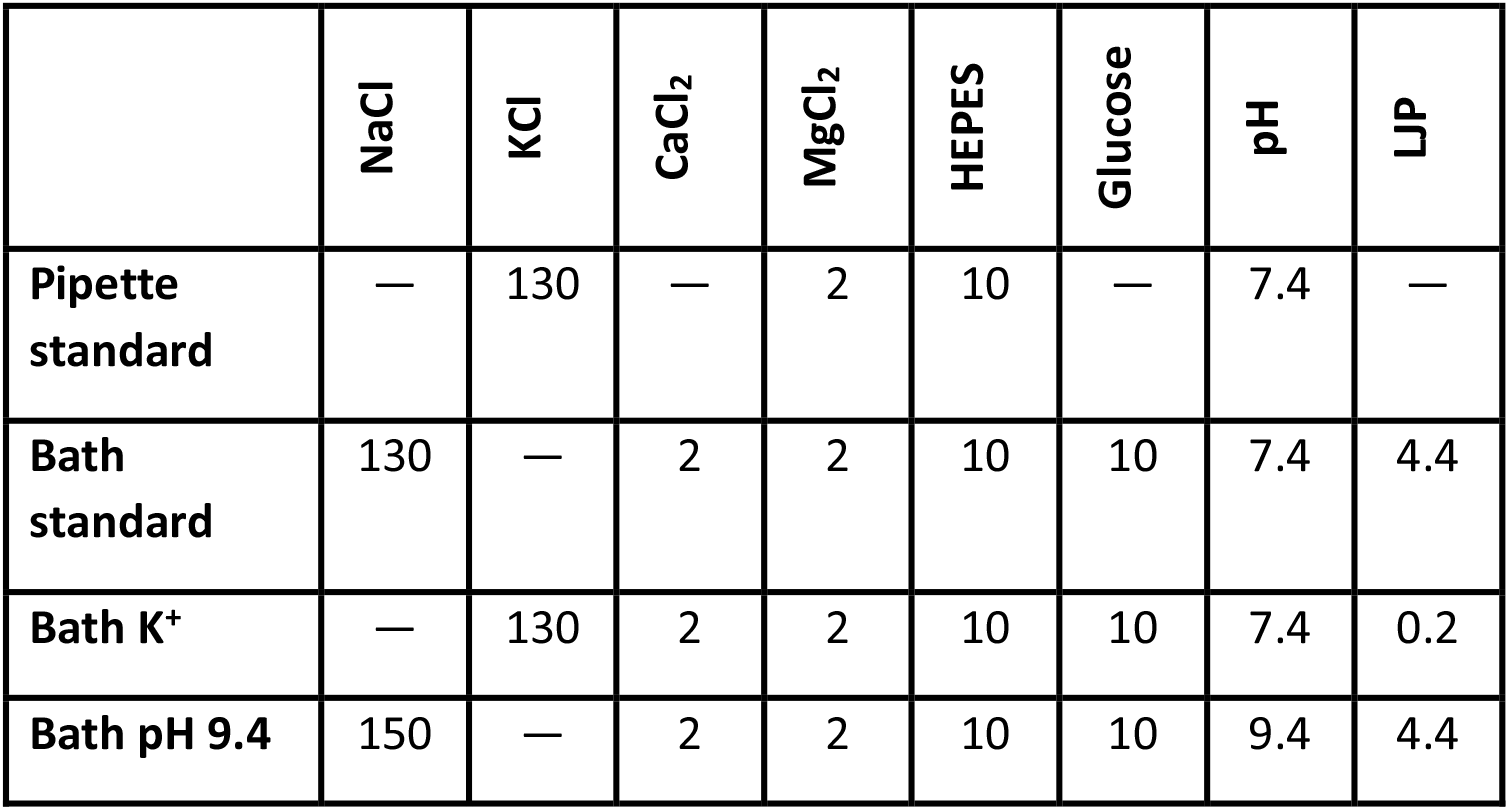
Solution compositions for whole-cell patch clamp recording. Abbreviations: HEPES, 4-(2-hydroxyethyl)-1-piperazineethanesulfonic acid; LJP, liquid junction potential. All concentrations are in mM.

**FIG S1.**
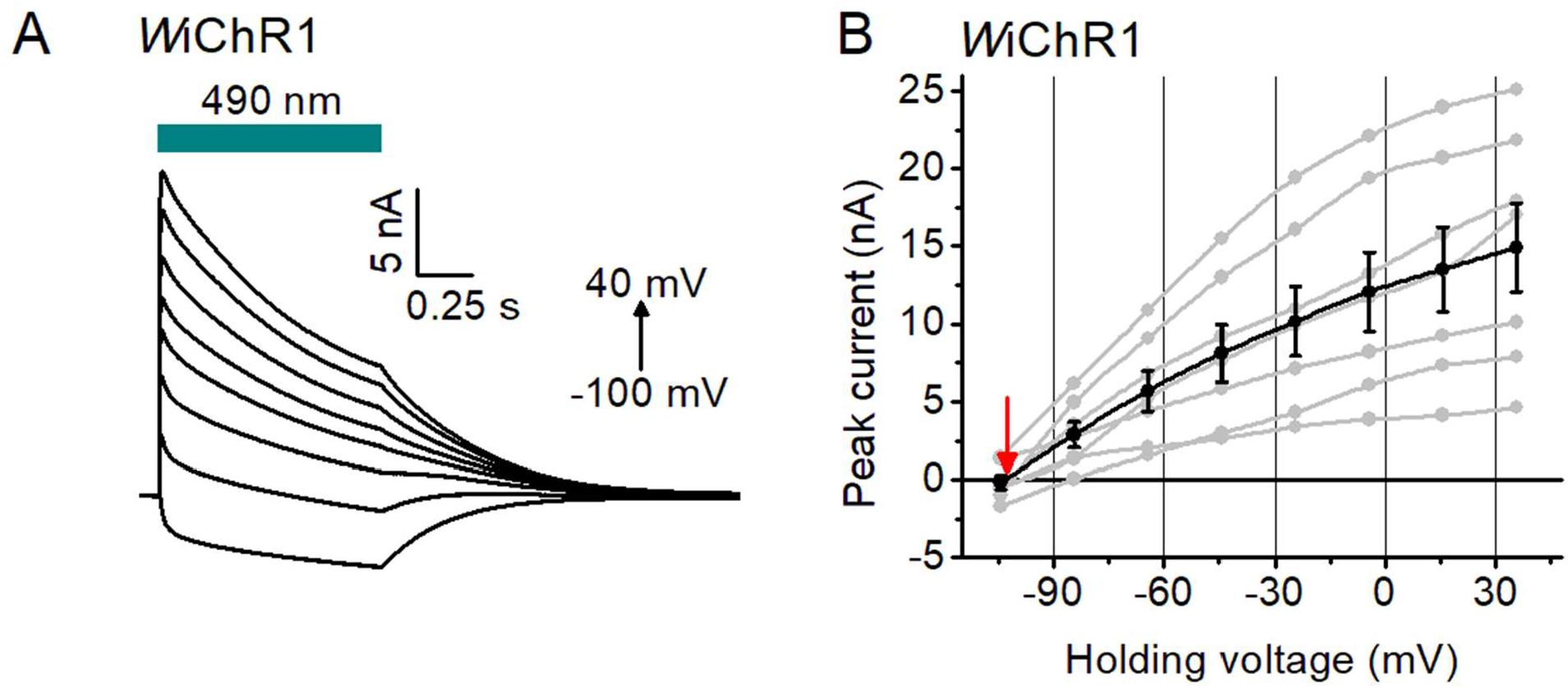
Electrophysiological characterization of *W*iChR1. (A) Series of photocurrent traces recorded upon incremental voltage with 130 mM Na^+^ in the bath and 130 mM K^+^ in the pipette. The duration of the light pulse is shown by the green bars. (C) The peak current-voltage relationships of *W*iChR1. Black, mean ± sem (n = 7 cells); grey, the data from individual cells. For two cells in which the photocurrent at the positive voltages exceeded the dynamic range of the amplifier (20 nA), the peak values were obtained by fitting of a sigmoidal (dose response) function to the data. The red arrow points to the reversal potential.

**FIG S2.**
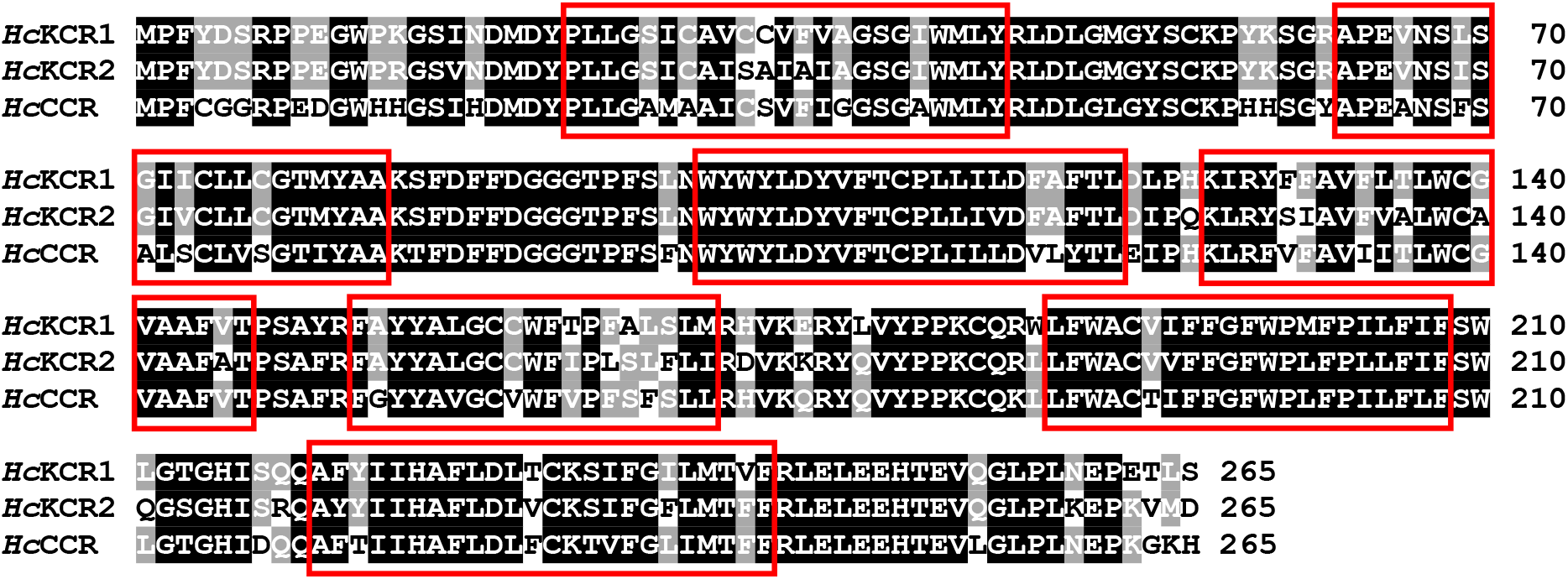
Protein alignment of 7TM domains of *H. catenoides* ChRs. The residues are shaded according to the degree of conservation. The numbers on the right are those of the last residue in each line. The red boxes show predicted transmembrane helices.

**FIG S3.**
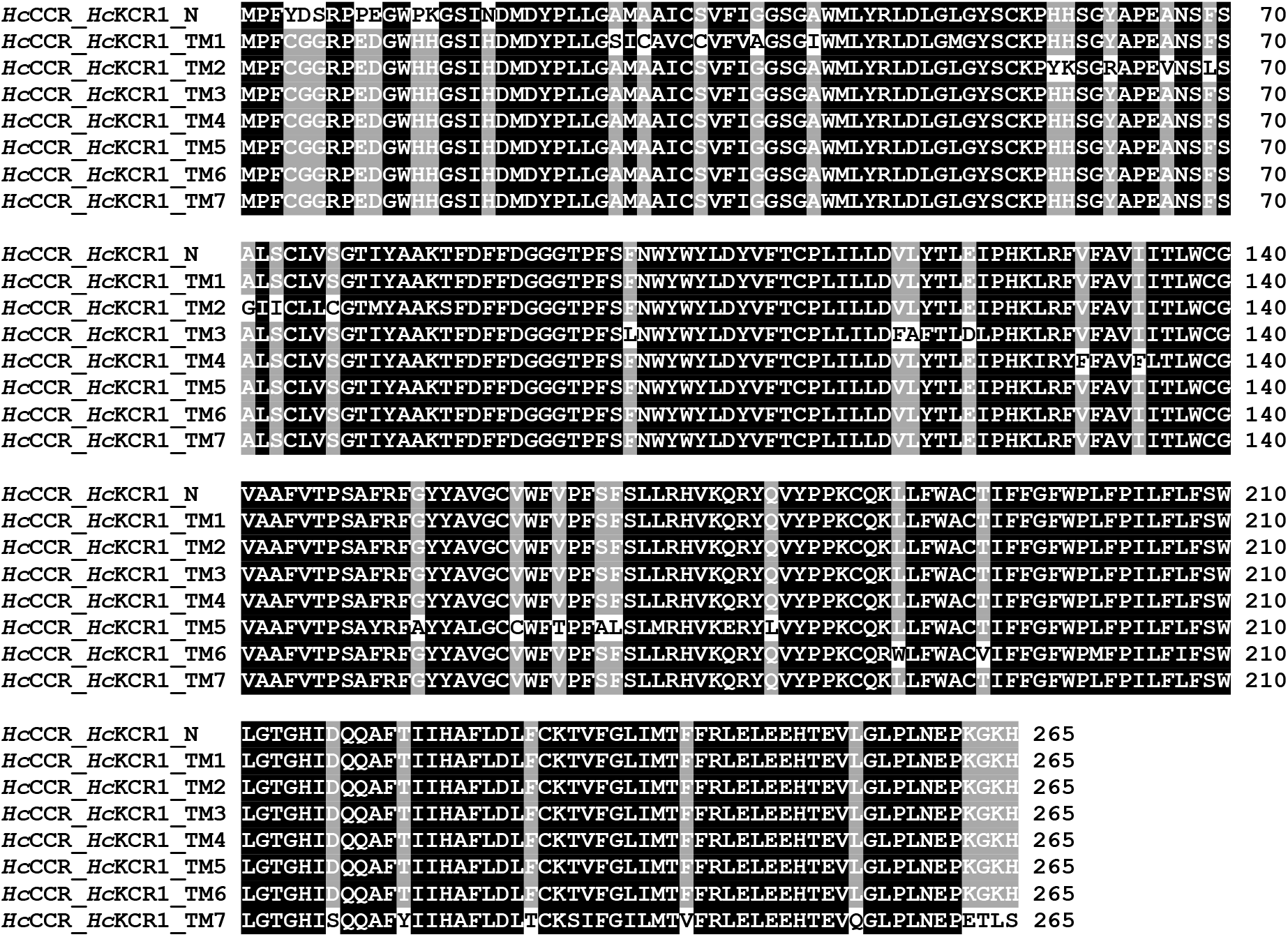
Protein alignment of the *Hc*KCR1_*Hc*CCR chimeras tested in this study. The residues are shaded according to the degree of conservation. The numbers on the right are those of the last residue in each line.

**FIG S4.**
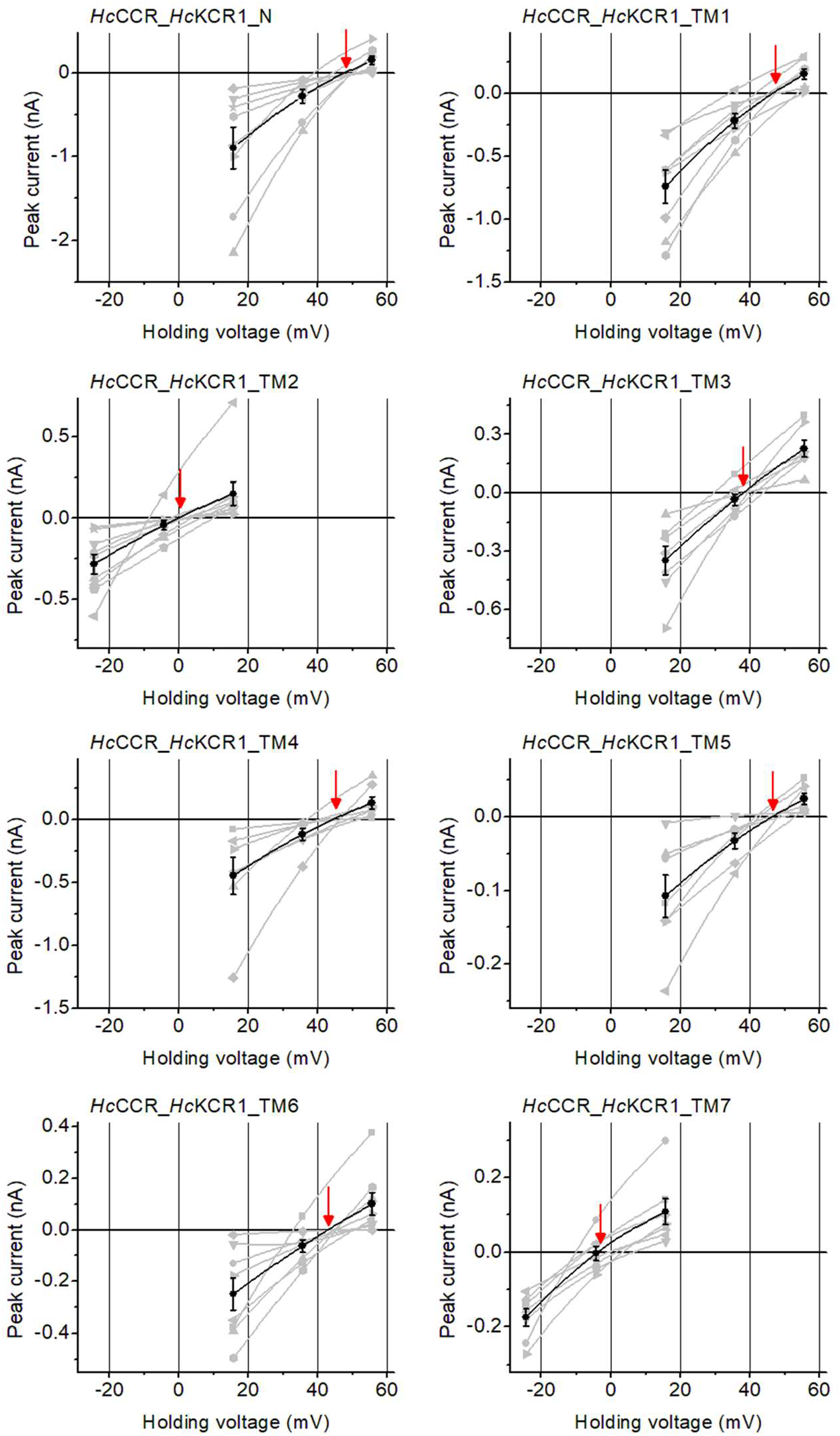
The peak current-voltage relationships of the *Hc*KCR1_*Hc*CCR chimeras. Black, mean ± sem (n = 7-10 cells); grey, the data from individual cells. The red arrows point to the reversal potentials.

**FIG S5.**
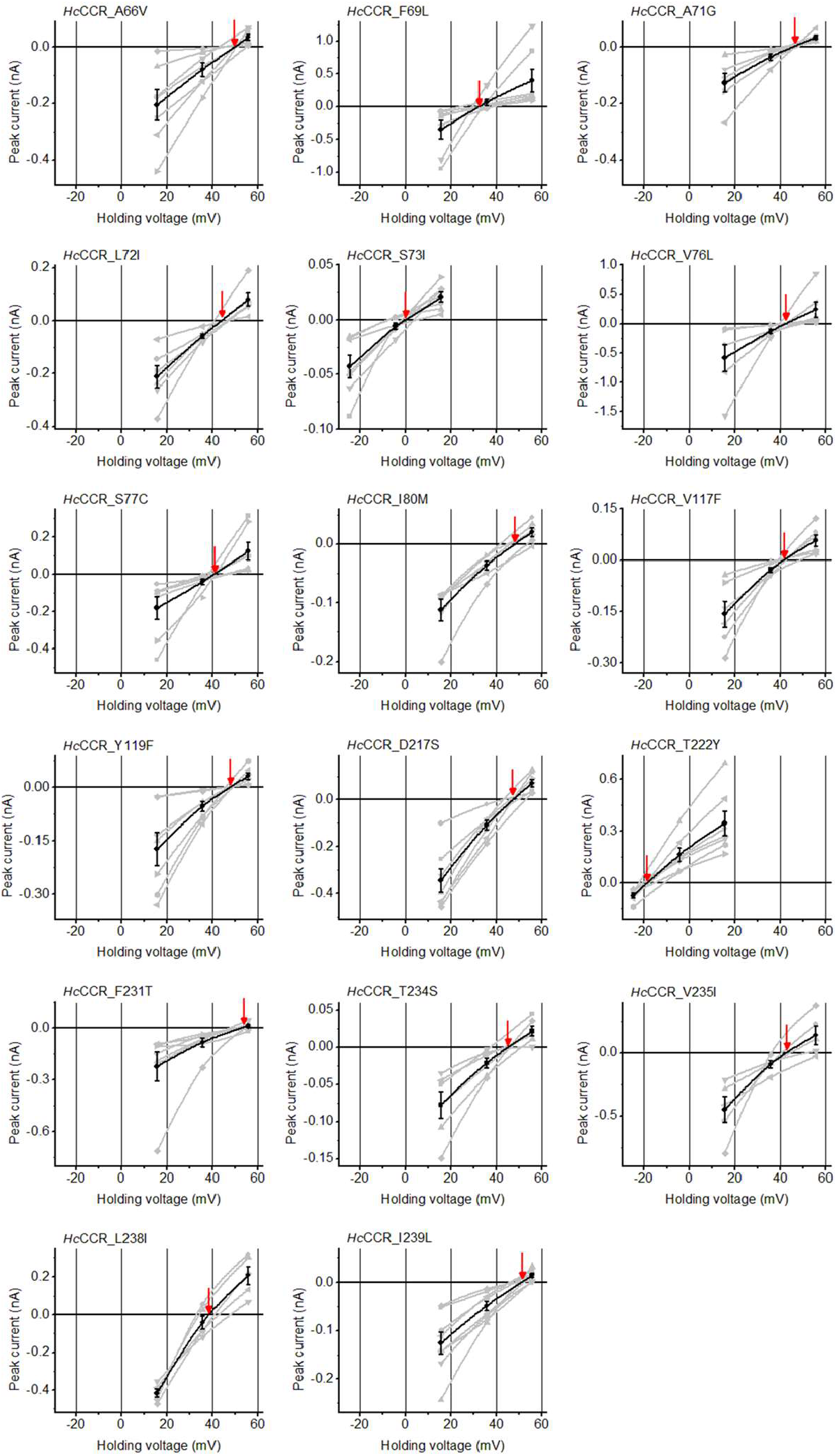
The peak current-voltage relationships of the *Hc*CCR mutants. Black, mean ± sem (n = 5-8 cells); grey, the data from individual cells. The red arrows point to the reversal potentials.

**FIG S6.**
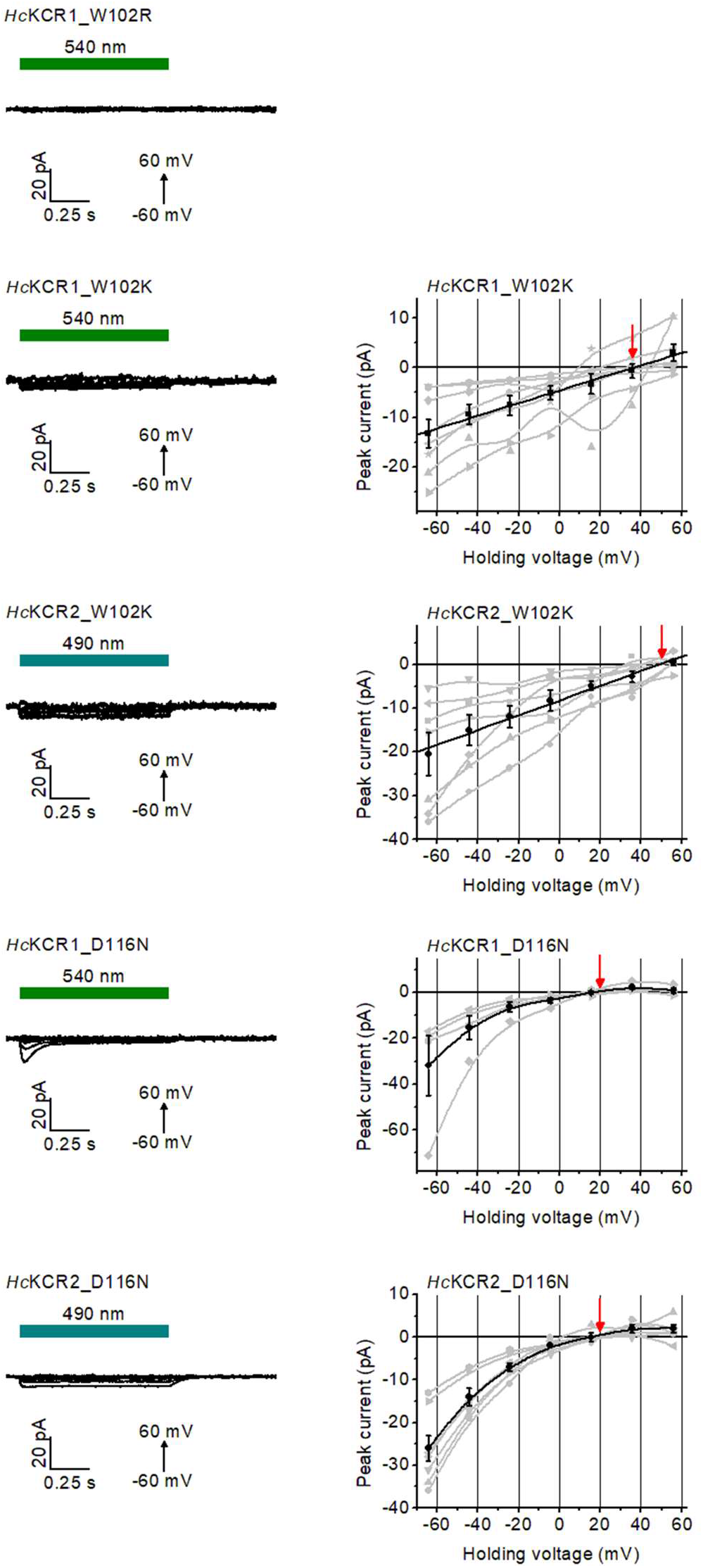
Characterization of Trp102 and Asp116 mutants of *Hc*KCR1 and *Hc*KCR2. Left, series of photocurrent traces recorded upon incremental voltage with 130 mM Na^+^ in the bath and 130 mM K^+^ in the pipette. The duration of the light pulse is shown by the green bars. Right, the peak current-voltage relationships. Black, mean ± sem (n = 7-8 cells); grey, the data from individual cells. The red arrows point to the reversal potentials.

**FIG S7.**
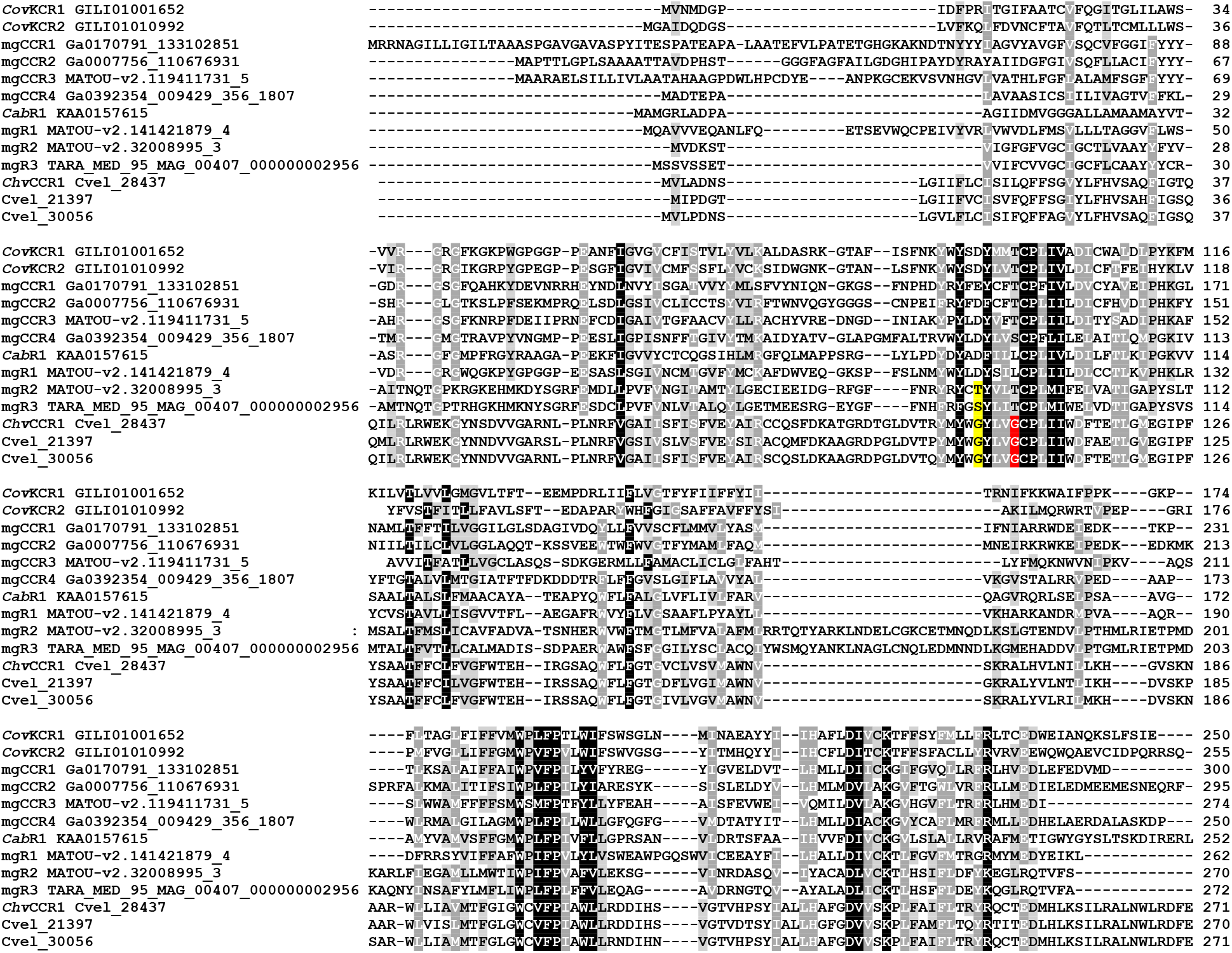
Protein alignment of 7TM domains of KCR homologs tested in this study. The residues are shaded according to the degree of conservation. In the *C. velia* sequences, the residues corresponding to Asp85 of bacteriorhodopsin are highlighted yellow, and the residues corresponding to Thr89 of bacteriorhodopsin, red. The numbers on the right are those of the last residue in each line.

**FIG S8.**
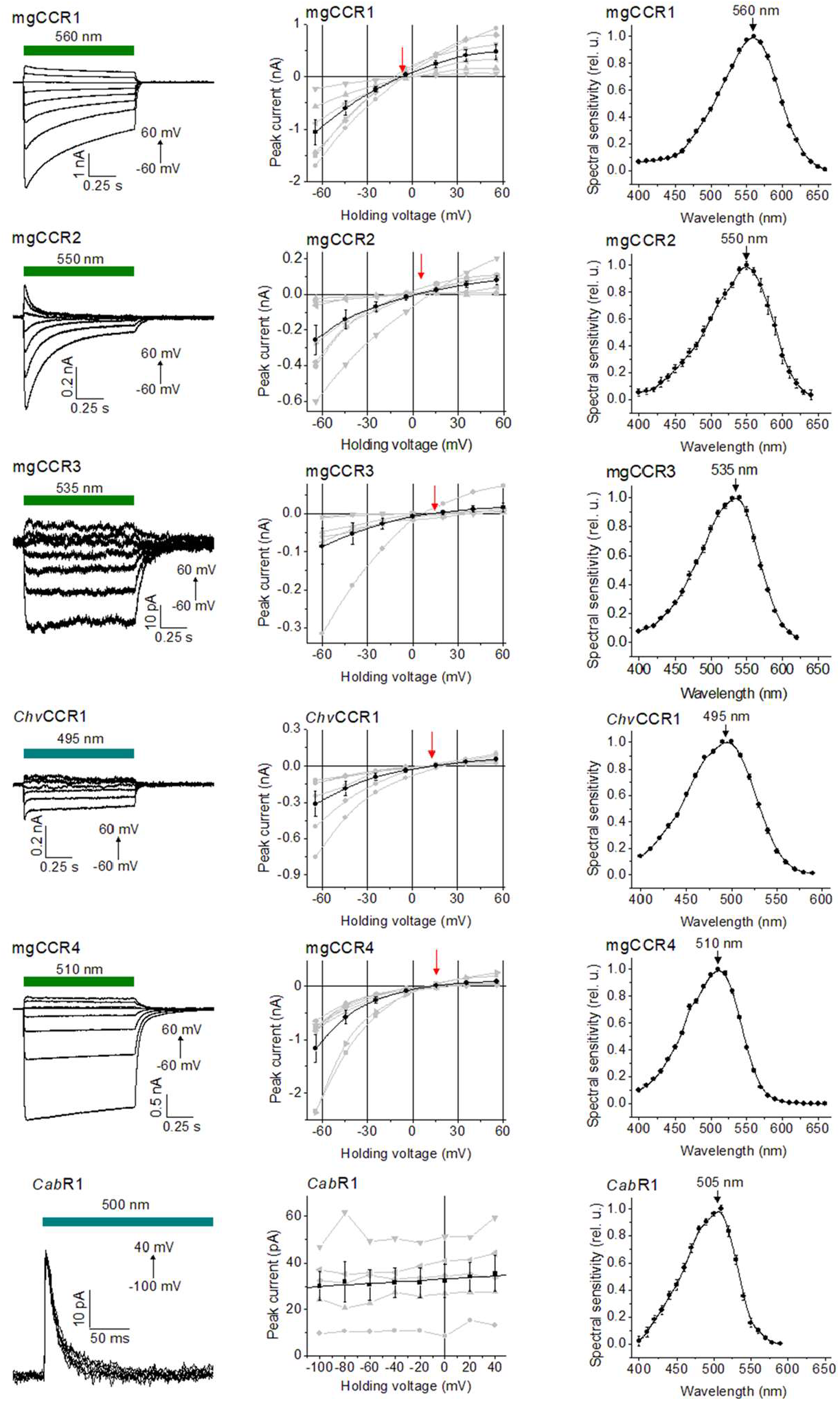
Characterization of KCR homologs with 130 mM Na^+^ in the bath. Left, series of photocurrent traces recorded upon incremental voltage. The duration of the light pulse is shown by the colored bars. Middle, the peak current-voltage relationships. Black, mean ± sem (n = 6-9 cells); grey, the data from individual cells. Right, the action spectra of the photocurrents (mean ± sem (n = 6-10 cells). The red arrows point to the reversal potentials.

**FIG S9.**
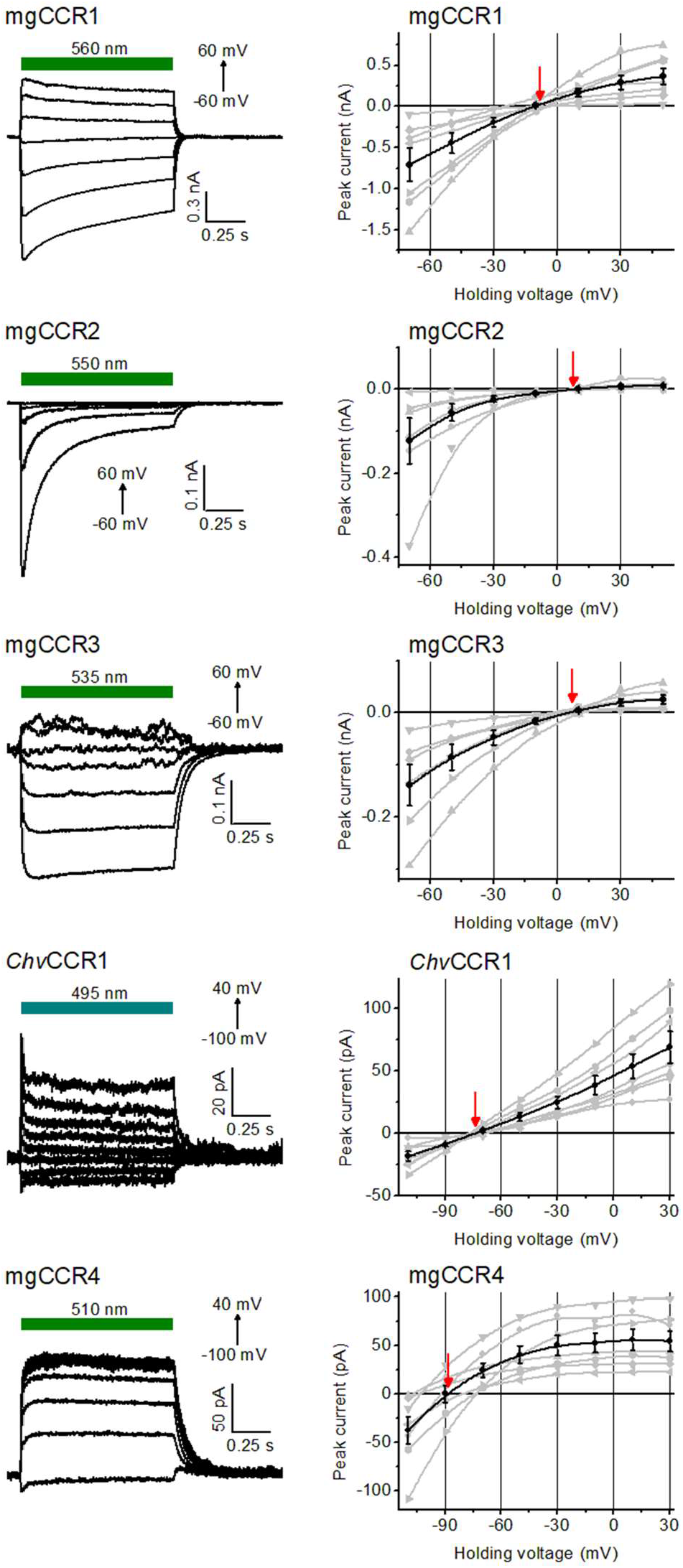
Characterization of KCR homologs with 130 mM NMDG^+^ in the bath. Left, series of photocurrent traces recorded upon incremental voltage. The duration of the light pulse is shown by the colored bars. Right, the peak current-voltage relationships. Black, mean ± sem (n = 6-7 cells); grey, the data from individual cells. The red arrows point to the reversal potentials.

**FIG S10.**
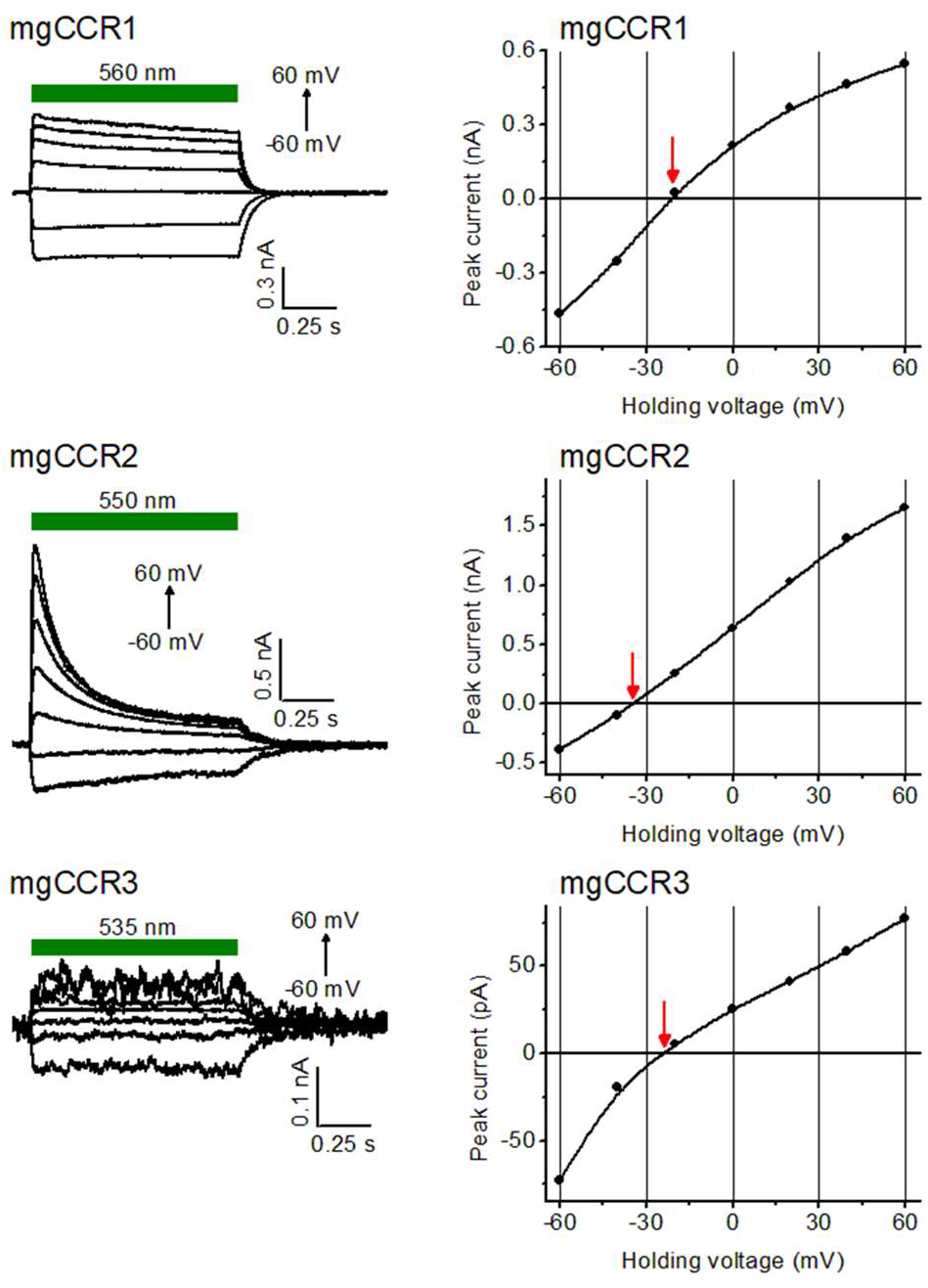
Characterization of KCR homologs with 130 mM K^+^ in the bath, pH 9.4. Left, series of photocurrent traces recorded upon incremental voltage. The duration of the light pulse is shown by the colored bars. Right, the corresponding peak current-voltage relationships. The red arrows point to the reversal potentials.

## REFERENCES

1. Govorunova EG, Sineshchekov OA, Li H, Wang Y, Brown LS, Palmateer A, Melkonian M, Cheng S, Carpenter E, Patterson J, Wong GKS, Spudich JL. 2021. Cation and anion channelrhodopsins: Sequence motifs and taxonomic distribution. MBio 12:e0165621.

2. Klapoetke NC, Murata Y, Kim SS, Pulver SR, Birdsey-Benson A, Cho YK, Morimoto, TK, Chuong AS, Carpenter EJ, Tian, Z, Wang J, Xie Y, Yan Z, Zhang Y, Chow BY, Surek B, Melkonian M, Jayaraman V, Constantine-Paton M, Wong GKS, Boyden ES. 2014. Independent optical excitation of distinct neural populations. Nat Methods 11:338–346.

3. Govorunova EG, Sineshchekov OA, Li H, Spudich JL. 2017. Microbial rhodopsins: Diversity, mechanisms, and optogenetic applications. Annu Rev Biochem 86:845–872.

4. Rozenberg A, Inoue K, Kandori H, Béjà O. 2021. Microbial rhodopsins: The last two decades. Annu Rev Microbiol 75:427–447.

5. Gordeliy V, Kovalev K, Bamberg E, Rodriguez-Valera F, Zinovev E, Zabelskii D, Alekseev A, Rosselli R, Gushchin I, Okhrimenko I. 2022. Microbial Rhodopsins. Methods Mol Biol 2501:1–52.

6. Sineshchekov OA, Jung K-H, Spudich JL. 2002. Two rhodopsins mediate phototaxis to low- and high-intensity light in Chlamydomonas reinhardtii. Proc Natl Acad Sci USA 99:8689–8694.

7. Govorunova EG, Sineshchekov OA, Liu X, Janz R, Spudich JL. 2015. Natural light-gated anion channels: A family of microbial rhodopsins for advanced optogenetics. Science 349:647–650.

8. Nagel G, Szellas T, Huhn W, Kateriya S, Adeishvili N, Berthold P, Ollig D, Hegemann P, Bamberg E. 2003. Channelrhodopsin-2, a directly light-gated cation-selective membrane channel. Proc Natl Acad Sci USA 100:13940–5.

9. Boyden ES, Zhang F, Bamberg E, Nagel G, Deisseroth K. 2005. Millisecond-timescale, genetically targeted optical control of neural activity. Nat Neurosci 8:1263–8.

10. Govorunova EG, Gou Y, Sineshchekov OA, Li H, Lu X, Wang Y, Brown LS, St-Pierre F, Xue M, Spudich JL. 2022. Kalium channelrhodopsins are natural light-gated potassium channels that mediate optogenetic inhibition. Nat Neurosci 25: 967–974.

11. MacKinnon R. 2003. Potassium channels. FEBS Lett 555:62–5.

12. Mironenko A, Zachariae U, de Groot BL, Kopec W. 2021. The persistent question of potassium channel permeation mechanisms. J Mol Biol 433:167002.

13. Sineshchekov OA, Govorunova EG, Li H, Spudich JL. 2017. Bacteriorhodopsin-like channelrhodopsins: Alternative mechanism for control of cation conductance. Proc Natl Acad Sci USA 114:E9512–E9519.

14. Kishi KE, Kim YS, Fukuda M, Inoue M, Kusakizako T, Wang PY, Ramakrishnan C, Byrne EFX, Thadhani E, Paggi JM, Matsui TE, Yamashita K, Nagata T, Konno M, Quirin S, Lo M, Benster T, Uemura T, Liu K, Shibata M, Nomura N, Iwata S, Nureki O, Dror RO, Inoue K, Deisseroth K, Kato HE. 2022. Structural basis for channel conduction in the pump-like channelrhodopsin ChRmine. Cell 185:672-689.e23.

15. Tucker K, Sridharan S, Adesnik H, Brohawn SG. 2022. Cryo-EM structures of the channelrhodopsin ChRmine in lipid nanodiscs. Nat Commun 13:4842.

16. Henderson R, Unwin P. 1975. Three-dimensional model of purple membrane obtained by electron microscopy. Nature 257:28–32.

17. Kato HE, Zhang F, Yizhar O, Ramakrishnan C, Nishizawa T, Hirata K, Ito J, Aita Y, Tsukazaki T, Hayashi S, Hegemann P, Maturana AD, Ishitani R, Deisseroth K, Nureki O. 2012. Crystal structure of the channelrhodopsin light-gated cation channel. Nature 482:369–374.

18. Li H, Huang CY, Govorunova EG, Schafer CT, Sineshchekov OA, Wang M, Zheng L, Spudich JL. 2019. Crystal structure of a natural light-gated anion channelrhodopsin. Elife 8:e41741.

19. Leonard G, Labarre A, Milner DS, Monier A, Soanes D, Wideman JG, Maguire F, Stevens S, Sain D, Grau-Bove X, Sebe-Pedros A, Stajich JE, Paszkiewicz K, Brown MW, Hall N, Wickstead B, Richards TA. 2018. Comparative genomic analysis of the ‘pseudofungus’ Hyphochytrium catenoides. Open Biol 8:170184.

20. Govorunova EG, Sineshchekov OA, Brown LS, Spudich JL. 2022. Biophysical characterization of light-gated ion channels using planar automated patch clamp. Front Mol Neurosci.

21. Hille B. 2001. Ion channels of excitable membranes. Sinauer Associates, Sunderland, MA.

22. Richards R, Dempski RE. 2012. Re-introduction of transmembrane serine residues reduce the minimum pore diameter of channelrhodopsin-2. PLoS One 7:e50018.

23. Vierock J, Peter E, Grimm C, Rozenberg A, Castro Scalise AG, Augustin S, Tanese D, Forget BC, Emiliani V, Béjà O, Hegemann P. 2022. WiChR, a highly potassium selective channelrhodopsin for low-light two-photon neuronal inhibition. bioRxiv doi:10.1101/2022.07.02.498568:2022.07.02.498568.

24. Otto H, Marti T, Holz M, Mogi T, Stern LJ, Engel F, Khorana HG, Heyn MP. 1990. Substitution of amino acids Asp-85, Asp-212, and Arg-82 in bacteriorhodopsin affects the proton release phase of the pump and the pK of the Schiff base. Proc Natl Acad Sci USA 87:1018–22.

25. Dolinsky TJ, Czodrowski P, Li H, Nielsen JE, Jensen JH, Klebe G, Baker NA. 2007. PDB2PQR: expanding and upgrading automated preparation of biomolecular structures for molecular simulations. Nucleic Acids Res 35:W522–5.

26. Tikhonenkov DV, Janouskovec J, Mylnikov AP, Mikhailov KV, Simdyanov TG, Aleoshin VV, Keeling PJ. 2014. Description of Colponema vietnamica sp.n. and Acavomonas peruviana n. gen. n. sp., two new alveolate phyla (Colponemidia nom. nov. and Acavomonidia nom. nov.) and their contributions to reconstructing the ancestral state of alveolates and eukaryotes. PLoS One 9:e95467.

27. Moore RB, Obornik M, Janouskovec J, Chrudimsky T, Vancova M, Green DH, Wright SW, Davies NW, Bolch CJ, Heimann K, Slapeta J, Hoegh-Guldberg O, Logsdon JM, Carter DA. 2008. A photosynthetic alveolate closely related to apicomplexan parasites. Nature 451:959–63.

28. Sineshchekov OA, Govorunova EG, Wang J, Li H, Spudich JL. 2013. Intramolecular proton transfer in channelrhodopsins. Biophys J 104:807–817.

29. Emiliani V, Entcheva E, Hedrich R, Hegemann P, Konrad KR, Lüscher C, Mahn M, Pan Z-H, Sims RR, Vierock J, Yizhar O. 2022. Optogenetics for light control of biological systems. Nature Reviews Methods Primers 2:55.

30. Sahel JA, Boulanger-Scemama E, Pagot C, Arleo A, Galluppi F, Martel JN, Esposti SD, Delaux A, de Saint Aubert JB, de Montleau C, Gutman E, Audo I, Duebel J, Picaud S, Dalkara D, Blouin L, Taiel M, Roska B. 2021. Partial recovery of visual function in a blind patient after optogenetic therapy. Nat Med 27:1223–1229.

31. Krause N, Engelhard C, Heberle J, Schlesinger R, Bittl R. 2013. Structural differences between the closed and open states of channelrhodopsin-2 as observed by EPR spectroscopy. FEBS Lett 587:3309–13.

32. Sattig T, Rickert C, Bamberg E, Steinhoff HJ, Bamann C. 2013. Light-induced movement of the transmembrane helix B in channelrhodopsin-2. Angew Chem Int Ed Engl 52:9705–8.

33. Müller M, Bamann C, Bamberg E, Kuhlbrandt W. 2015. Light-induced helix movements in channelrhodopsin-2. J Mol Biol 427:341–9.

34. Li H, Huang C-Y, Govorunova EG, Sineshchekov OA, Yi A, Rothschild KJ, Wang M, Zheng L, Spudich JL. 2021. The crystal structure of bromide-bound GtACR1 reveals a pre-activated state in the transmembrane anion tunnel. Elife 10:e65903.

35. Brown LS, Gat Y, Sheves M, Yamazaki Y, Maeda A, Needleman R, Lanyi JK. 1994. The retinal Schiff base-counterion complex of bacteriorhodopsin: changed geometry during the photocycle is a cause of proton transfer to aspartate 85. Biochemistry 33:12001–11.

36. Wietek J, Wiegert JS, Adeishvili N, Schneider F, Watanabe H, Tsunoda SP, Vogt A, Elstner M, Oertner TG, Hegemann P. 2014. Conversion of channelrhodopsin into a light-gated chloride channel. Science 344:409–412.

37. Vogt A, Silapetere A, Grimm C, Heiser F, Ancina Moller M, Hegemann P. 2019. Engineered passive potassium conductance in the KR2 sodium pump. Biophys J 116:1941–1951.

38. Sineshchekov OA, Govorunova EG, Li H, Wang Y, Melkonian M, Wong GK-S, Brown LS, Spudich JL. 2020. Conductance mechanisms of rapidly desensitizing cation channelrhodopsins from cryptophyte algae. mBio 11:e00657–20.

39. Kumpf RA, Dougherty DA. 1993. A mechanism for ion selectivity in potassium channels: computational studies of cation-pi interactions. Science 261:1708–10.

40. Otto H, Marti T, Holz M, Mogi T, Lindau M, Khorana HG, Heyn MP. 1989. Aspartic acid-96 is the internal proton donor in the reprotonation of the Schiff base of bacteriorhodopsin. Proc Natl Acad Sci USA 86:9228–32.

41. Nagel G, Ollig D, Fuhrmann M, Kateriya S, Musti AM, Bamberg E, Hegemann P. 2002. Channelrhodopsin-1: a light-gated proton channel in green algae. Science 296:2395–8.

42. Rozenberg A, Kaczmarczyk I, Matzov D, Vierock J, Nagata T, Sugiura M, Katayama K, Kawasaki Y, Konno M, Nagasaka Y, Aoyama M, Das I, Pahima E, Church J, Adam S, Borin VA, Chazan A, Augustin S, Wietek J, Dine J, Peleg Y, Kawanabe A, Fujiwara Y, Yizhar O, Sheves M, Schapiro I, Furutani Y, Kandori H, Inoue K, Hegemann P, Béjà O, Shalev-Benami M. 2022. Rhodopsin-bestrophin fusion proteins from unicellular algae form gigantic pentameric ion channels. Nat Struct Mol Biol 29:592–603.

43. Villar E, Vannier T, Vernette C, Lescot M, Cuenca M, Alexandre A, Bachelerie P, Rosnet T, Pelletier E, Sunagawa S, Hingamp P. 2018. The Ocean Gene Atlas: exploring the biogeography of plankton genes online. Nucleic Acids Res 46:W289–W295.

44. Vernette C, Lecubin J, Sanchez P, Tara Oceans C, Sunagawa S, Delmont TO, Acinas SG, Pelletier E, Hingamp P, Lescot M. 2022. The Ocean Gene Atlas v2.0: online exploration of the biogeography and phylogeny of plankton genes. Nucleic Acids Res doi:10.1093/nar/gkac420:DOI: 10.1093/nar/gkac420.

45. Carradec Q, Pelletier E, Da Silva C, Alberti A, Seeleuthner Y, Blanc-Mathieu R, Lima-Mendez G, Rocha F, Tirichine L, Labadie K, Kirilovsky A, Bertrand A, Engelen S, Madoui MA, Meheust R, Poulain J, Romac S, Richter DJ, Yoshikawa G, Dimier C, Kandels-Lewis S, Picheral M, Searson S, Tara Oceans C, Jaillon O, Aury JM, Karsenti E, Sullivan MB, Sunagawa S, Bork P, Not F, Hingamp P, Raes J, Guidi L, Ogata H, de Vargas C, Iudicone D, Bowler C, Wincker P. 2018. A global ocean atlas of eukaryotic genes. Nat Commun 9:373.

46. Delmont TO, Gaia M, Hinsinger DD, Fremont P, Vanni C, Guerra AF, Eren AM, Kourlaiev A, d’Agata L, Clayssen Q, Villar E, Labadie K, Cruaud C, Poulain J, Da Silva C, Wessner M, Noel B, Aury J-M, de Vargas C, Bowler C, Karsenti E, Pelletier E, Wincker P, Jaillon O. 2021. Functional repertoire convergence of distantly related eukaryotic plankton lineages revealed by genome-resolved metagenomics. bioRxiv doi:10.1101/2020.10.15.341214:2020.10.15.341214.

47. Chen IA, Chu K, Palaniappan K, Ratner A, Huang J, Huntemann M, Hajek P, Ritter S, Varghese N, Seshadri R, Roux S, Woyke T, Eloe-Fadrosh EA, Ivanova NN, Kyrpides NC. 2021. The IMG/M data management and analysis system v.6.0: new tools and advanced capabilities. Nucleic Acids Res 49:D751–D763.

48. Grigoriev IV, Hayes RD, Calhoun S, Kamel B, Wang A, Ahrendt S, Dusheyko S, Nikitin R, Mondo SJ, Salamov A, Shabalov I, Kuo A. 2021. PhycoCosm, a comparative algal genomics resource. Nucleic Acids Res 49:D1004–D1011.

49. Woo YH, Ansari H, Otto TD, Klinger CM, Kolisko M, Michalek J, Saxena A, Shanmugam D, Tayyrov A, Veluchamy A, Ali S, Bernal A, del Campo J, Cihlar J, Flegontov P, Gornik SG, Hajduskova E, Horak A, Janouskovec J, Katris NJ, Mast FD, Miranda-Saavedra D, Mourier T, Naeem R, Nair M, Panigrahi AK, Rawlings ND, Padron-Regalado E, Ramaprasad A, Samad N, Tomcala A, Wilkes J, Neafsey DE, Doerig C, Bowler C, Keeling PJ, Roos DS, Dacks JB, Templeton TJ, Waller RF, Lukes J, Obornik M, Pain A. 2015. Chromerid genomes reveal the evolutionary path from photosynthetic algae to obligate intracellular parasites. Elife 4:e06974.

50. Hallgren J, Tsirigos KD, Pedersen MD, Almagro Armenteros JJ, Marcatili P, Nielsen H, Krogh A, Winther O. 2022. DeepTMHMM predicts alpha and beta transmembrane proteins using deep neural networks. bioRxiv doi:10.1101/2022.04.08.487609:2022.04.08.487609.

51. Minh BQ, Schmidt HA, Chernomor O, Schrempf D, Woodhams MD, von Haeseler A, Lanfear R. 2020. IQ-TREE 2: New models and efficient methods for phylogenetic inference in the genomic era. Mol Biol Evol 37:1530–1534.

52. Mirdita M, Schutze K, Moriwaki Y, Heo L, Ovchinnikov S, Steinegger M. 2022. ColabFold: making protein folding accessible to all. Nat Methods 19:679–682.

53. Furuse M, Tamogami J, Hosaka T, Kikukawa T, Shinya N, Hato M, Ohsawa N, Kim SY, Jung KH, Demura M, Miyauchi S, Kamo N, Shimono K, Kimura-Someya T, Yokoyama S, Shirouzu M. 2015. Structural basis for the slow photocycle and late proton release in Acetabularia rhodopsin I from the marine plant Acetabularia acetabulum. Acta Crystallogr D Biol Crystallogr 71:2203–16.

54. Brooks BR, Bruccoleri RE, Olafson BD, States DJ, Swaminathan S, Karplus M. 1983. CHARMM: a program for macromolecular energy, minimization, and dynamics calculations. J Comput Chem 4:187–217.

55. MacKerell AD, Bashford D, Bellott M, Dunbrack RL, Evanseck JD, Field MJ, Fischer S, Gao J, Guo H, Ha S, Joseph-McCarthy D, Kuchnir L, Kuczera K, Lau FT, Mattos C, Michnick S, Ngo T, Nguyen DT, Prodhom B, Reiher WE, Roux B, Schlenkrich M, Smith JC, Stote R, Straub J, Watanabe M, Wiorkiewicz-Kuczera J, Yin D, Karplus M. 1998. All-atom empirical potential for molecular modeling and dynamics studies of proteins. J Phys Chem B 102:3586–616.

56. Mackerell AD, Jr., Feig M, Brooks CL, 3rd. 2004. Extending the treatment of backbone energetics in protein force fields: limitations of gas-phase quantum mechanics in reproducing protein conformational distributions in molecular dynamics simulations. J Comput Chem 25:1400–15.

57. Jorgensen WL, Chandrasekhar J, Madura JD, Impey RW, Klein ML. 1983. Comparison of simple potential functions for simulating liquid water. J Chem Phys 79:926–935.

58. Nina M, Roux B, Smith JC. 1995. Functional interactions in bacteriorhodopsin: a theoretical analysis of retinal hydrogen bonding with water. Biophys J 68:25–39.

59. Tajkhorshid E, Baudry J, Schulten K, Suhai S. 2000. Molecular dynamics study of the nature and origin of retinal’s twisted structure in bacteriorhodopsin. Biophys J 78:683–693.

60. Gruia AD, Bondar AN, Smith JC, Fischer S. 2005. Mechanism of a molecular valve in the halorhodopsin chloride pump. Structure 13:617–27.

